# Bayesian-Steered Structure Prediction of Mechanical Biomolecules Using Twisted Diffusion

**DOI:** 10.64898/2026.05.11.724187

**Authors:** Colin Klaus, Marcos Sotomayor

## Abstract

Deep learning approaches have revolutionized protein structure prediction. These tools are trained using experimental data and recapitulate reported conformations, but there is great interest in predicting conformations that may be functionally relevant although experimentally underrepresented. Since many modern structure prediction tools use generative artificial intelligence diffusion models, we reframe the search for alternative molecular conformations as that of sampling from a diffusion distribution conditioned using any arbitrary Bayesian likelihood. We implement a twisted diffusion sampler in Boltz-2 to sample this conditioned distribution and demonstrate the utility of this approach, which does not require any additional training of the neural network, by implementing a diffusion analog of steered molecular dynamics simulations applied to mechanical systems. We can reproduce predicted stretched states of fragments of DNA, the muscle protein titin, and the inner-ear protocadherin-15 protein, as well as open states of the MscL ion channel consistent with experimental results. We expect that steered structure predictions will help sample underrepresented and non-equilibrium conformations for many macromolecular systems.

## Introduction

The field of protein structure prediction and design has been revolutionized by deep learning tools (1) such as AlphaFold (2, 3), ESMFold (4), OpenFold (5), Boltz (6, 7), and others (8–13). These tools often use neural networks trained on experimental biomolecular structures deposited and accumulated over decades in the protein data bank (PDB) (14), which was the foundation for their success. At the same time, structures in the PDB may underrepresent functionally relevant conformations that are difficult to experimentally characterize, such as stretched states of mechanical proteins and open states of mechanosensitive channels. Consequently, models trained on the PDB data inherently favor generating structures similar to the training set, even if these models have the capacity to predict physically accurate alternate conformations. There is also great interest in customizing predictions to exhibit additional properties required by a user. This challenge has motivated the development of *ad hoc* methods for increasing prediction diversity. Within AlphaFold2, one of the most prevalent and empirically successful approaches has been to manipulate the multiple-sequence-alignment (MSA) (15–19). Other notable approaches have been to incorporate distance constraints by either retraining or fine-tuning the neural network (20–22), to optimize custom-loss functions post-training (23), or to rescale the latent pair representation (24). AlphaFold3’s structure prediction module amounted to a paradigm shift compared to its predecessor. While AlphaFold2 (2) used an innovative residue gas approach and assembled the backbone using rigid body motions over per residue frames of reference, AlphaFold3 (3) employed a diffusion model (25–28) directly over the atomic coordinates. Since during training diffusion models learn a probability distribution over a discretized path space, the implicit statistical potential over that path space serves as a natural loss function for optimizing structure predictions subject to additional user defined constraints. More generally, Bayesian approaches provide a technical framework for generating ensembles of structure predictions that sample this implicit statistical potential after adding additional restraint potentials that enforce user constraints (29, 30). These techniques can often be administered without any additional training of the underlying neural network (25, 28, 31–33).

Here we customize Boltz-2’s, an open-source version of AlphaFold3, to predict biomolecular structures that satisfy arbitrary properties set by a user. Formally, we sample the diffusion distribution learned by Boltz-2 after conditioning it with a Bayesian likelihood that encodes the user-desired constraints. We approximately sample these conditional distributions by implementing twisted diffusion (32), a hybrid sampling technique that combines strategies introduced for classifier guidance (28, 31) and sequential Monte Carlo (SMC) (30, 34, 35) inside Boltz-2 (Figure 1). As a concrete application, we take normally distributed likelihoods whose means are arbitrary functions of the atomic coordinates called collective variables (colvars). In analogy with molecular dynamics simulations (36–39), conditioning the diffusion distribution with these likelihoods amounts to harmonically restraining these colvars to specified values with a spring constant that is determined by the likelihood’s standard deviation. We benchmark this strategy on some of the first biomolecules stretched using single-molecule force spectroscopy, such as DNA and a fragment of the muscle protein titin (40–50). We also consider fragments of the inner-ear force-conveying protocadherin-15 (PCDH15) protein and the heterodimeric interface between PCDH15 and cadherin-23 (CDH23) essential for vertebrate hearing and balance (51–58). For all these systems, we perform a diffusion analog of steered molecular dynamics (SMD) simulations (59–68) using end-to-end distance as the colvar. In this manner, we quantify what Boltz-2 predicts for the stretched and unfolded states of these biomolecules. Lastly, we apply this strategy to the pentameric mechanosensitive channel of large conductance (MscL) from *Mycobacterium tuberculosis* (69). Although natively, Boltz-2 only predicts the closed state of MscL, we find that conditioning Boltz-2 with an appropriate colvar generates structural predictions for the open state transmembrane region that are consistent with models predicted using different approaches (70–75).

**Figure 1:**
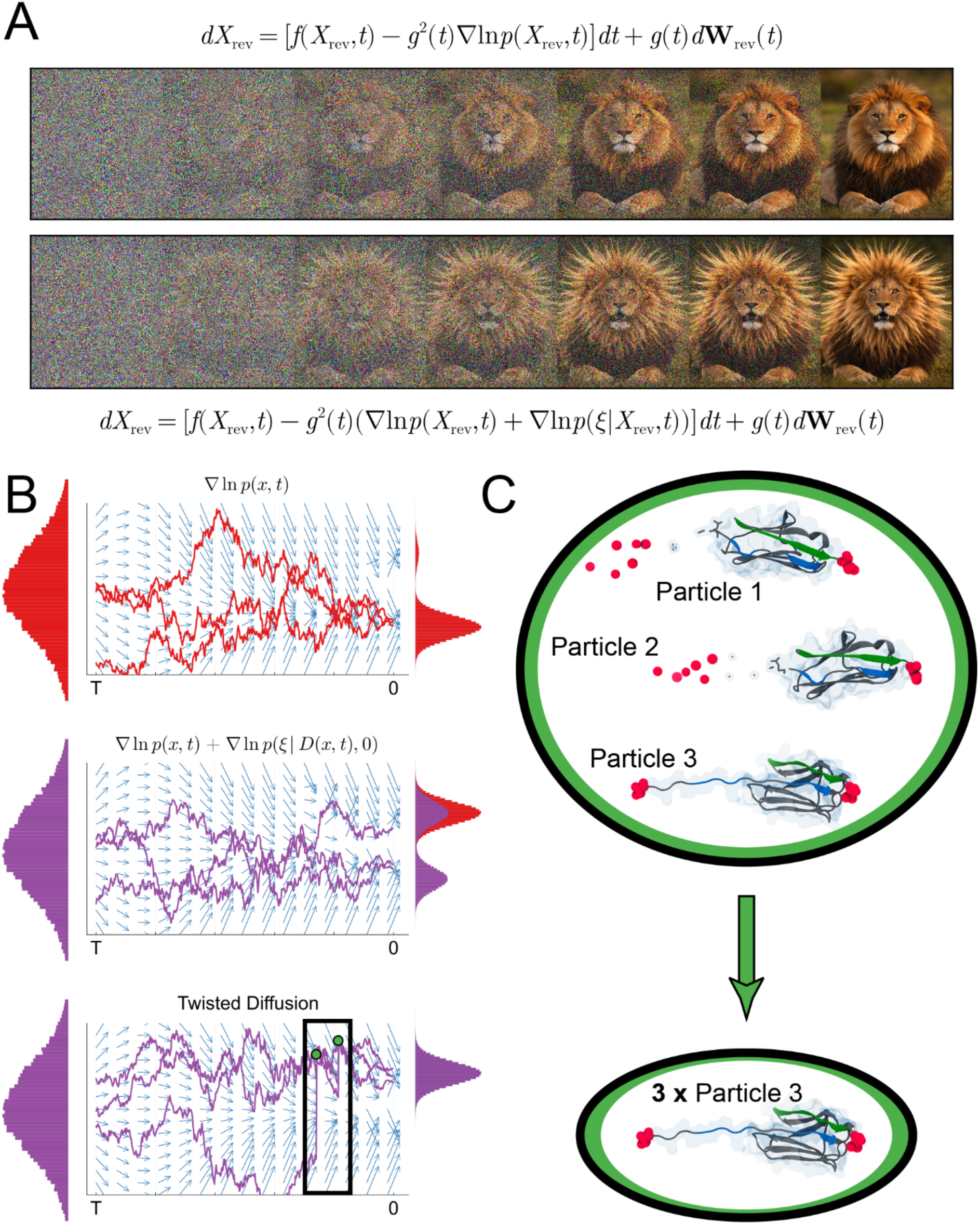
Illustration of inference time steering and twisted diffusion. *A*. Diffusion models can be steered post-training to produce samples with additional properties exploiting that they are score based. An illustration of unsteered diffusion producing an image of a lion (top) using the unconditional score 𝛻 𝑙𝑛 𝑝 (𝑥, 𝑡). An illustration of steered diffusion producing an image of a lion whose mane has been extended (bottom) using the conditional score 𝛻 𝑙𝑛 𝑝 (𝑥, 𝑡 | 𝜉) and Bayes’ formula. Lion images are intended as a schematic and were not generated through a full twisted diffusion pipeline (see **Methods**). *B*. Samples with rare properties are hardly generated by an unsteered diffusion process (top). The diffusion process can be steered to better sample rare properties by approximating the true but intractable Bayes’ term with a similar term built from the denoiser (middle). Twisted diffusion is a sequential Monte Carlo scheme that asymptotically corrects the error due to approximating the true Bayes’ term with a proxy. Multiple trajectories are launched simultaneously, and poorer trajectories can leap to more promising regions of sample space found by the other trajectories for better exploration (bottom). *C*. For structure prediction, the implication of the sequential Monte Carlo framework is that intermediate structures are scored according to how well they satisfy the desired properties and resemble the original data distribution. Poorer scoring samples have a lower chance of surviving to the next iteration of diffusion, while higher scoring samples have a higher chance of being replicated for future refinement.

## Results

### Bayesian-steered Boltz-2

We created a Bayesian-steered version of Boltz-2 that can accept custom constraints set by a user and produce structure predictions that are compatible with those constraints. We first explain the broad conceptual changes made to the standard Boltz-2 approach (see **Methods** and **Appendices I and II** for technical details).

By default, Boltz-2 takes an input sequence of amino acids, nucleic acids, or ligands and uses that information to guide a reverse diffusion process that transforms pure Gaussian noise into a biomolecular structure (7). The process begins at a mathematically formal, and nonphysical, step 𝑡_𝑛_by initializing the position of all atoms with independent samples of a Gaussian distribution with standard deviation 𝑡_𝑛_. These coordinates are stored in the variable 𝑋_𝑡_𝑛__. In the variance exploding (VE) elucidated diffusion model (EDM) framework (76), which AlphaFold3 and Boltz-2 adapt (3, 7), the reverse diffusion alternates between adding new but diminishingly small Gaussian noise to the atom positions and denoising steps that walk the noisy coordinates towards a denoised prediction made with the current coordinates. More specifically, let 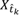 denote the atomic coordinates at step *t̂*_𝑘_ after adding new Gaussian noise with standard deviation 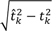 to the coordinates 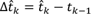. The less noisy atomic coordinates at step 𝑡_𝑘−1_ are obtained using

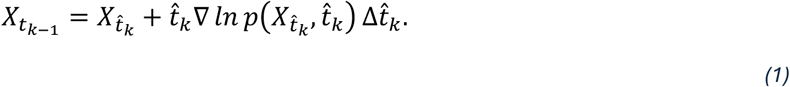

where 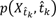 is the marginal probability density of the diffusion process at time *t̂*_𝑘_ and evaluated at 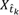. This process iterates until reaching the ideal 𝑡_0_ = 0 where the prediction 𝑋_0_finally contains no noise. The key term in equation *(1)* is the so-called *score* ∇ 𝑙𝑛 𝑝 (𝑥, 𝑡). During training, the VE EDM learns a denoiser that approximates the best mean-square-error predictor of the original, noiseless coordinates given the current noisy coordinates. This denoiser, 𝐷(𝑥, 𝑡), and the score are related through denoising score matching (26, 28, 77) which gives

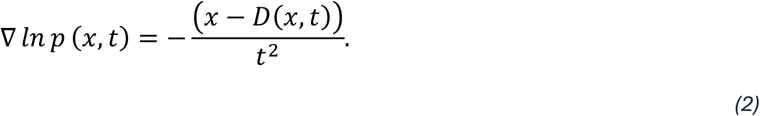

Substituting equation *(1)* into *(2)*, we obtain how the denoiser guides the VE EDM’s denoising process

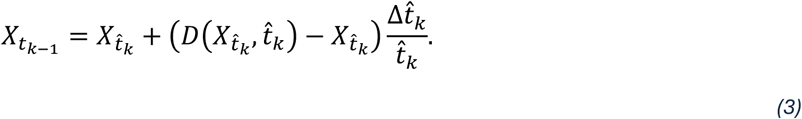

In our Bayesian-steered Boltz-2 code the denoising process described by equation *(3)* is first modified by biasing the denoiser’s predictions to better satisfy the desired user constraints. These constraints are input by the user into our code as a restraint potential 𝑈(𝑥; ξ) whose value depends on a colvar ξ. The associated Bayesian likelihood is defined as 𝑝(ξ|𝑥_0_, 0) ∝ 𝑒𝑥𝑝(−𝑈(𝑥_0_; ξ)) so that equality holds up to normalization. In the language of Bayesian inference, adding this restraint potential into the implicit statistical potential over structure trajectories originally learned by Boltz-2 is equivalent to conditioning Boltz-2’s distribution on diffusion trajectories with 𝑝(ξ|𝑥_0_, 0). Under conditioning, the score transforms according to Bayes’ formula (28) which combined with equation *(2)* implies 𝐷(𝑥, 𝑡; ξ) = 𝐷(𝑥, 𝑡) + 𝑡^2^∇ 𝑙𝑛 𝑝 (ξ|𝑥, 𝑡), defining the denoiser for the conditional diffusion process. Although theoretically exact, the ∇ 𝑙𝑛 𝑝 (ξ|𝑥, 𝑡) term is intractable at diffusion steps 𝑡 > 0, but the denoiser is used to define an inexact approximation (32)

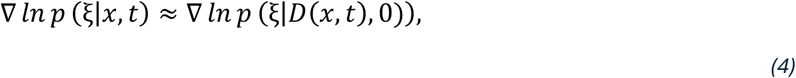

where we differentiate through the denoiser according to chain rule. Based on these arguments, in our Bayesian-steered Boltz-2 code equation *(3)* becomes

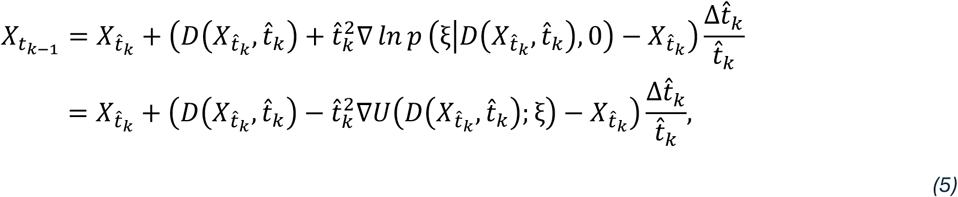

This modification of the denoising dynamics leads to a mechanism for proposing structures that better satisfy the user constraints. However, due to the inexact approximation described by equation *(4)*, the code does not yet sample from Boltz-2’s original structure distribution tilted by the weight factor 𝑒𝑥𝑝(−𝑈(𝑥_0_; ξ)/ ∝ 𝑝(ξ|𝑥_0_, 0). This tilted distribution is also called the posterior distribution. To correct that error, we implement twisted diffusion (32) which is an example of Feynman-Kac steering (33). A population of noisy structures, also called Feynman-Kac particles, undergo reverse diffusion and at intermediate steps are weighted and then are replicated or killed in an asymptotically exact manner (30, 34, 35). These weighting and *resampling* procedures explicitly depend on the user prescribed Bayesian likelihood defining the constraint, equivalent to the restraint potential 𝑈(𝑥; ξ). In the limit of increasingly many particles, *i.e.*, simultaneous structure predictions, the terminal population at step 𝑡 = 0 tends to an exact approximation of the true posterior.

To use our Bayesian-steered Boltz-2, the user first defines the colvar ξ and restraint potential 𝑈(𝑥; ξ). These terms are defined in a new `cxolvars.py` python file and may be built using any function that pytorch can automatically differentiate. The user also specifies the number of diffusion steps and the number of Feynman-Kac particles. Several other hyperparameters for improving performance, such as tempering coefficients for the statistical energy and gradient steering, are also adjustable. We additionally provide a Jupyter notebook (`Boltz2AtomsIds.ipynb)` that reconstructs Boltz-2’s internal listing of residues by atoms which is used to make atom selections for defining colvars. In the second half of the notebook, a `connected_atom_index.pt` pytorch tensor is exported. This tensor lists additional atom indices, which in our applications were indices for the 3’ oxygen and the 5’ phosphate atoms forming the DNA phosphodiester bond or for the C and N backbone atoms forming the peptide bond within a protein chain. These atom indices are then included within Boltz-2’s `ConnectionsPotential` class to penalize these atoms from exceeding physical bond distances under stretching. More detailed documentation for running our customized Boltz-2 is available on github (see also **Methods**, **Appendix I: Methods**).

### Applications of Bayesian-steered Boltz-2

To test our Bayesian-steered Boltz-2 predictions of stretched and unfolded states of biomolecules, we began with simple but also historically significant model systems (DNA and titin) followed with inner-ear PCDH15 protein fragments and the CDH23 and PCDH15 protein complex with ligands (Ca^2+^). We end with predictions for the pentameric MscL channel. Each system poses challenges of increasing complexity. We expect to see untwisting of the DNA double helix and unfolding of titin, unbending and unrolling for PCDH15 fragments with some differences caused by ligands, and unbinding for the CDH23 and PCDH15 complex. For MscL, we expect to be able to see assembly of the pentameric complex in open conformations that are not present in the training data set. We did unsteered predictions for each system and applied four progressively larger end-to-end distance harmonic restraints for all systems except MscL, for which we tested two open pore sizes. Five independent predictions (𝑛 = 5) each using 100 Feynman- Kac particles were obtained for each harmonic distance restraint or pore size, with additional parameters described in Methods. The resulting highest probability particle, *i.e.*, prediction, was analyzed for each case. Additionally, 500 default predictions using Boltz-2’s original settings were also performed for MscL. Sequences (*STable 1*), but no templates or additional structural information, were used for all predictions. No training or re-training of the Boltz-2 neural network was required either.

### Predictions for end-to-end stretching of a DNA fragment show elongation and untwisting

We first applied a harmonic colvar bias to the terminal base pairs (bp) of a 12-bp DNA double helix fragment with a sequence matching the initial portion of λ-DNA stretched in early single-molecule force spectroscopy experiments (*40*) (*STable 1*).

In unsteered predictions, the DNA fragment was predicted to form a double helix spanning 32.8 ± 2.0 Å as measured by the distance between the terminal base pairs’ center of masses (Figure 2A, SFigure 1A). DNA strands were correctly arranged in an antiparallel fashion and the 5’ to 3’ end-to-end distance was 35.1 ± 3.2 Å. The observed distances were shorter than the expected length of 40.8 Å (12 × 3.4 Å). Harmonically restraining the terminal bps to 45 Å led to an elongated but still intact double helix spanning 42.7 ± 1.0 Å (Figure 2A, SFigure 1B). At a harmonic restraint of 55 Å, the double helix spanned 52.0 ± 0.9 Å. Continuing to a restraint of 65 Å, the double helix spanned 60.3 ± 0.7 Å. Several predictions showed partial untwisting of the helix with one case showing complete untwisting. Predictions without untwisting still showed large deformations of the helical turn (Figure 2A, SFigure 1D). Finally, a harmonic restraint of 75 Å resulted in end-to-end distances of 68.1 ± 1.0 Å. This case showed similar trends as what was observed at 65 Å, but the majority of predictions showed untwisting. In one case, shredding was predicted (Figure 2A, SFigure 1E). Interestingly, some predictions showed shredding involving one to three atoms, including three predictions for unsteered DNA. Overall, our *in-silico* findings are consistent with the fact that DNA is an extensible biomolecule that can be stretched up to twice its length (40, 42–45). The predicted conformations are also consistent with results from prior SMD simulations (78–80).

**Figure 2:**
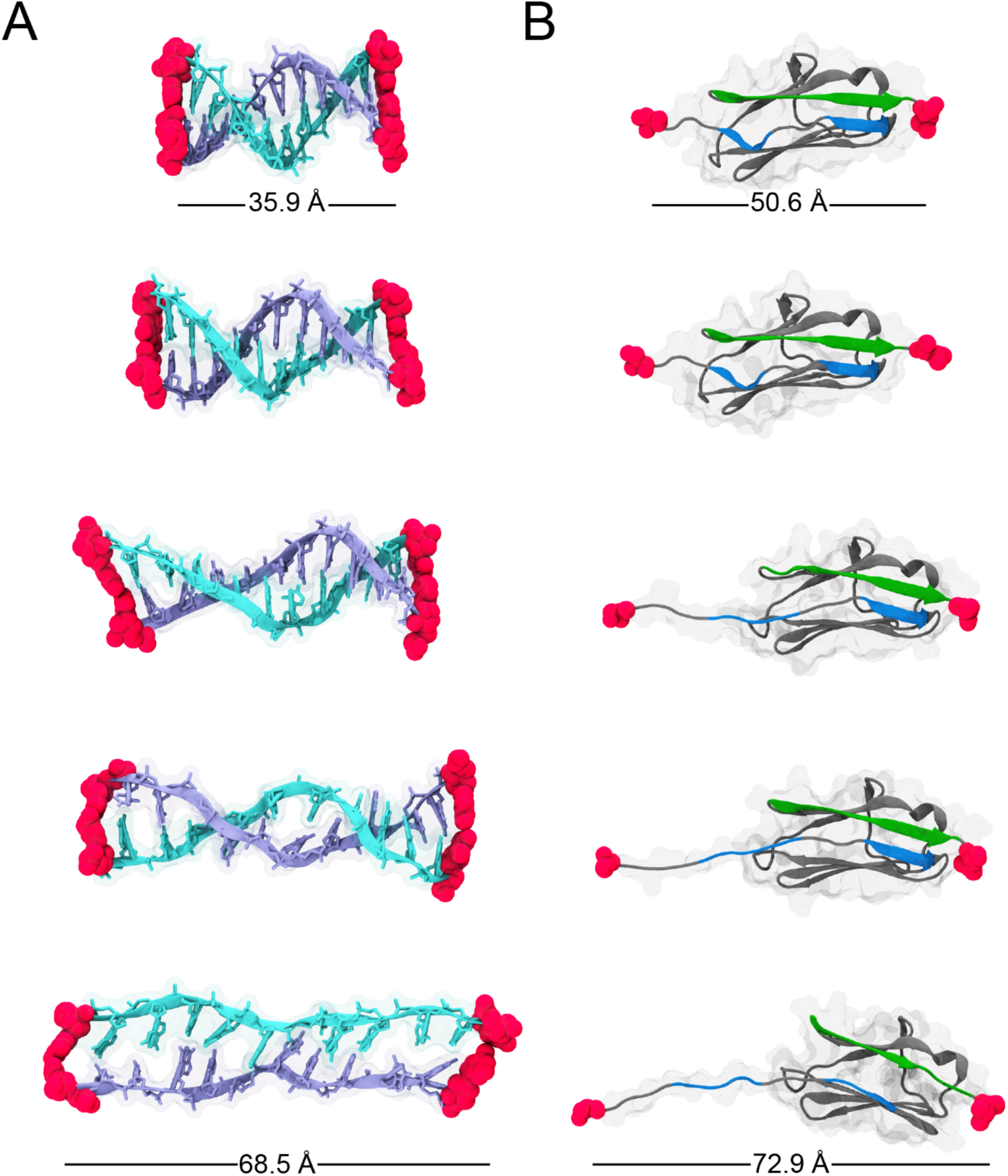
Bayesian-steered Boltz-2 predictions for DNA and titin IG1 at varying end-to-end distances. *A*. Stretching predictions for a 12-base pair DNA double helix. The colvar was defined as the distance between centers of mass of each pair of terminal bases. The colvar was harmonically restrained from 35 Å – 75 Å. DNA strands are shown in cyan and ice blue licorice and cartoon representations with transparent molecular surfaces. Atoms used to define the colvar are shown as red spheres. *B*. Stretching predictions for a titin I91 domain (also known as I27). The colvar was defined as the distance between centers of mass of the N- and C-terminal residues. The colvar was harmonically restrained over a range of 55 Å – 75 Å. Protein is shown in gray cartoon representation with a transparent molecular surface. Terminal β strands A, A’, and G are shown in blue and green. Targeted atoms are shown as red spheres. Figure panels are also shown in SFigure 1 and SFigure 2.

### Predictions for end-to-end stretching of a titin domain show sequential unfolding of β strands

Our second system consisted of a single titin IG domain (I91, also known as I27). Titin is a very large protein, and its passive elasticity is important for muscle function (81). The I91 domain (91 amino acids, *STable 1*), extensively studied using both single-molecule force spectroscopy experiments and SMD simulations (46–50, 61), is mechanically stable requiring large forces to unfold (> 200 pN at room temperature and 100 μm/s) (50). This domain has a β-sandwich fold with two sheets formed by β-strands ABED and A’GFC. The large energetic cost of shearing out β-strands during stretching is thought to explain its mechanical resilience. We obtained titin predictions using an end-to-end distance colvar (N- and C-terminal residues) and compared these to its available experimental structure (PDB: 1TIT), obtained from nuclear magnetic resonance (NMR) experiments (82), and to predictions of its stretched and unfolded states from SMD simulations (41, 48, 49, 61).

In unsteered predictions, the titin I91 domain was folded with a backbone root mean squared deviation (RMSD) of 1.3 ± 0.1 Å compared to the deposited NMR structure (PDB: 1TIT) (82). The end-to-end distance between its N- and C-terminal residues was 46.9 ± 3.0 Å (Figure 2B, SFigure 2A). Where their sequences overlapped, that end-to-end distance (sequence-overlap end-to-end distance) of the NMR structure was 43.2 Å and the Boltz-2 predictions showed 43.4 ± 0.6 Å. Harmonically restraining the end-to-end distance to 60 Å led to a stretching of the N- and C-terminal residues to 55.7 ± 1.9 Å (Figure 2B, SFigure 2B). In three of these predictions, β strands A and B detached from each other (SFigure 2B). Increasing the harmonic restraint to 65 Å, led to a stretching of 62.9 ± 0.7 Å. Three of the predictions showed *shredding* at the N-terminal residue but the other two showed predictions that were qualitatively similar to those obtained at the previous restraint distance (Figure 2B, SFigure 2C). Harmonically restraining the end-to-end distance to 70 Å led to stretching of 66.8 ± 0.6 Å. Here only two predictions showed shredding at the N-terminal end and were otherwise similar to what was observed using the previous restraint distance (SFigure 2D). Finally, increasing the harmonic restraint to 75 Å led to a stretching of the end-to-end distance to 71.3 ± 1.1 Å. Across the three predictions that did not show shredding, all showed unbinding of the A β strand and one showed unbinding of the A’ β strand from the G β strand (Figure 2B, SFigure 2E).

The conformations predicted by Bayesian-steered Boltz-2 models of stretched titin I91 are consistent with SMD simulations that show detachment of the A and B β strands followed by detachment of the A’ and G β strands (41). However, across all our steered Boltz-2 predictions, we did not observe extraction of β strand G as seen in some SMD simulations (41). This may occur at larger extensions not yet tested with our approach. Interestingly, in some steered predictions at the maximum harmonic restraint tested we observed formation of a β-sheet involving β-strands A’ and B, rather than A’ and G. This suggests that our Bayesian-steered Boltz-2 predictions may capture distinct stretched I91 folds that are difficult to access in fast SMD simulations.

### Predictions for end-to-end stretching of a bent PCDH15 fragment show straightening before unfolding

After validating our approach using DNA and titin, we tested our approach on larger biomolecules consisting of fragments of Ca^2+^-dependent mechanical proteins essential for inner-ear function. These involved fragments of the *tip link*’s molecular constituents, PCDH15 and CDH23, which mechanically gate the ion channel involved in hearing and balance (51, 58, 83). The first of these inner-ear-related systems was the monomeric PCDH15 fragment including extracellular cadherin (EC) repeats 8, 9, and 10 (EC8-10, 322 amino acids, *STable 1*). Cadherin EC repeats, with seven β strands forming a Greek-key motif, are about 100 amino acids each and are similar in fold and sequence, but not identical to each other (84). The EC-EC linkers typically feature up to three bound Ca^2+^ ions that rigidify the structure and favor a straight conformation as observed for the PCDH15 EC8-9 linker (85). In contrast, experimental structures have revealed that the PCDH15 EC9-10 linker is unique as it lacks Ca^2+^-binding sites and it can be found in bent and straight conformations (85, 86). The bent conformation features a small 3_10_-helix connecting EC9 and EC10 (85). We used an end-to-end colvar (N- and C-terminal residues) to obtain structural predictions and compared them to available crystal structures of the same fragment, stretched conformations from SMD (85), and straightened conformations obtained using cryo-EM (86).

We predicted structures for PCDH15 EC8-10 both with and without Ca^2+^. When present, Ca^2+^ occupied the expected linker region at the EC8-9 linker in all but three cases. Otherwise our findings were similar in both cases. In unsteered predictions, PCDH15 EC8-10 with Ca^2+^ exhibited a backbone RMSD of 7.1 ± 1.7 Å compared to the available crystal structure (PDB: 4XHZ) (85), which also had Ca^2+^. Predictions without Ca^2+^ exhibited a backbone RMSD of 6.9 ± 2.3 Å compared to that same crystal structure. These predictions showed the characteristic bent conformation of EC9 and EC10 (85) (Figure 3AB, SFigure 3A, SFigure 4A). However, the RMSD values were heavily influenced by differences between the EC9-10 bending angles for predictions and the experimental structure. RMSD values obtained when just aligning EC8-9 were 3.3 ± 0.6 Å and 4.1 ± 0.4 Å with and without Ca^2+^, respectively. In contrast RMSD values obtained when aligning EC9-10 were 5.4 ± 2.3 Å and 5.3 ± 2.5 Å with and without Ca^2+^ respectively. Interestingly, a comparison with the EC9-10 fragment from another similar PCDH15 structure (PDB: 6EET (51)) resulted in smaller RMSD values (4.7 ± 2.2 Å and 5.0 ± 2.5 Å with and without Ca^2+^, respectively). The predicted end-to-end distances were 101.4 ± 7.3 Å and 100.3 ± 7.3 Å and the sequence-overlap end-to-end distances were 111.6 Å for the experimental structure and 101.4 ± 7.4 Å and 100.1 ± 7.4 Å for predictions with and without Ca^2+^, respectively. As with RMSD values, sequence-overlap distance discrepancies reflected differences in angles. These results indicate that unsteered predictions are accurate for individual EC repeats, capture the straight conformation of the EC8-9 fragment with Ca^2+^, predict only straight conformations for the EC8-9 fragment without Ca^2+^, and capture the bent conformation of EC9-10 albeit with angles that differ from those observed in crystal structures (51, 85).

**Figure 3:**
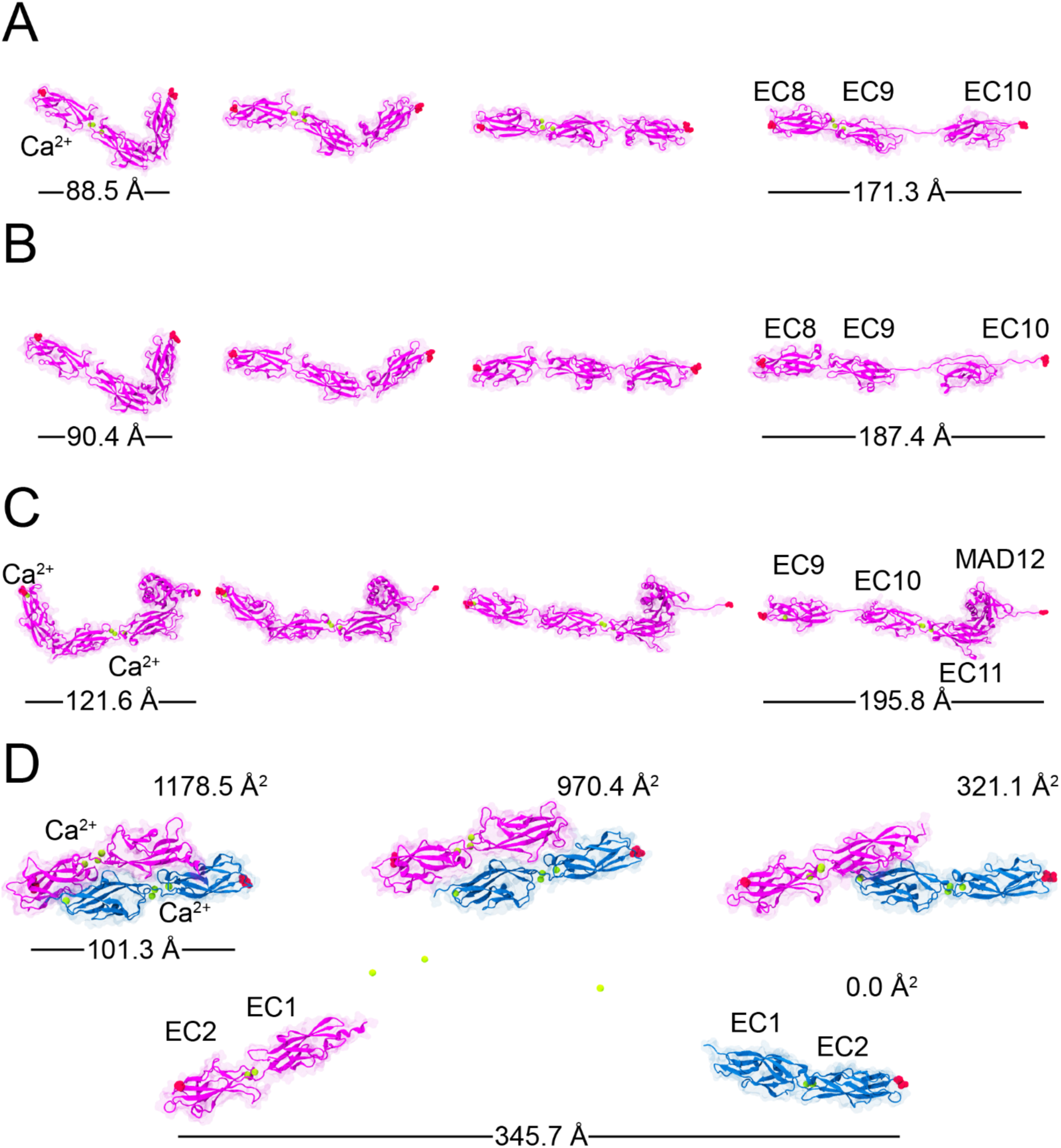
Bayesian-steered Boltz-2 predictions for selected inner-ear-related protein fragments at varying end-to-end distances. *A*-*B*. Stretching predictions for PCDH15 EC8-10 with and without Ca^2+^ at canonical binding sites, respectively. PCDH15 is shown in magenta cartoon and transparent surface representations. Ca^2+^ ions are shown as green spheres. Targeted atoms are shown as red spheres. *C*. Stretching predictions for PCDH15 EC9-MAD12 with Ca^2+^ at canonical binding sites. Shown as in *A*. For *A* to *C* predictions the colvar was defined as the distance between centers of mass of the N- and C-terminal residues and was harmonically restrained over a range of 110 Å – 210 Å. *D*. Stretching predictions for the dimeric CDH23 EC1-2 and PCDH15 EC1-2 handshake with Ca^2+^ at canonical binding sites. Proteins are shown as in *A* with CDH23 in blue. The colvar was harmonically restrained over a range of 100 Å – 350 Å for C-terminal residues. Buried surface area (BSA) is also shown in units of Å^2^. Figure panels are also shown in SFigure 3, SFigure 4, SFigure 5, and SFigure 6.

Bayesian-steered predictions for PCDH15 EC8-10 using an end-to-end harmonic restraint of 130 Å demonstrated distances of 124.3 ± 0.9 Å and 127.0 ± 4.6 Å with and without Ca^2+^ with EC9-10 bending still visible in all but one case (Figure 3A, B, SFigure 3B, SFigure 4B). Two predictions without Ca^2+^ showed unfolding at the small 3_10_-helix in the EC9-10 linker region. Upon applying an end-to-end harmonic restraint of 160 Å, the stretched distances were 141.7 ± 1.0 Å and 142.7 ± 1.2 Å with and without Ca^2+^, respectively. At this threshold, predictions showed straightening of the EC9-10 fragment with only unfolding at the 3_10_-helix (Figure 3A, B, SFigure 3C, SFigure 4C). One prediction showed a misplaced Ca^2+^ ion. After applying an end-to-end harmonic restraint of 210 Å, the stretched distances were 179.3 ± 6.1 Å and 179.4 ± 4.2 Å with and without Ca^2+^. All predictions with and without Ca^2+^ showed unfolding of the 3_10_-helix and extraction of β-strands. Four of the five predictions with Ca^2+^ showed unfolding near the EC9-10 linker region. One prediction showed unfolding near the EC8-9 linker region, with Ca^2+^ ions misplaced. Four predictions with Ca^2+^ also showed slight to modest unfolding at the C-terminal end (Figure 3A, SFigure 3D). All five of the predictions without Ca^2+^ showed unfolding near their EC9-10 linker region as well as unfolding at the C-terminal end (Figure 3B, SFigure 4D). Applying a final harmonic restraint of 260 Å, the end-to-end distances predicted were 205.9 ± 2.7 Å and 207.0 ± 3.8 Å with and without Ca^2+^. All predictions with and without Ca^2+^ showed unfolding of the EC9-10 3_10_- helix and extraction of β-strands. All five of the predictions with Ca^2+^ showed unfolding near the EC9-10 linker region while four also showed unfolding at the C-terminal end and one had the EC8-9 linker extended with Ca^2+^ ions misplaced. All five of the predictions without Ca^2+^ showed unfolding near the EC9- 10 linker region. In one case EC9 unfolded and split with part of it detaching from the rest. The EC9-10 linker showed more extension than the EC8-9 linker. All of these predictions without Ca^2+^ again showed unfolding at the C-terminal end (Figure 3B, SFigure 4E).

Overall, our Bayesian-steered Boltz-2 predictions for the PCDH15 EC8-10 fragment showed that the bent EC9-10 linker was the first to extend and only when imposing larger stretching did a β strand unfold. These results are consistent with spontaneously occurring straightened conformations observed in cryo-EM (86) and with SMD simulations of PCDH15 EC8-10 showing EC9-10 straightening before unfolding (85), suggesting also that Bayesian-steered Boltz-2 can capture states associated with tertiary structure elasticity of proteins (41).

### Predictions for end-to-end stretching of a PCDH15 fragment including a membrane adjacent domain

The second inner-ear related system was a similar fragment of the monomeric PCDH15 now including its membrane adjacent domain (MAD). The PCDH15 EC9-MAD12 fragment (448 amino acids, *STable 1*) includes the Ca^2+^-free EC9-10 linker, a canonical EC10-EC11 linker with up to three Ca^2+^ ions, and MAD12 with a ferredoxin-like fold that includes two α helices and four β strands (βαββαβ) (86, 87). Unlike the EC repeats that connect in series and have N- and C-termini at opposite ends, MAD12 tucks against EC11 and its N and C termini are adjacent to each other, with the C-terminal end followed by a post-MAD12 α helix that connects to PCDH15’s transmembrane domain. SMD simulations of monomeric PCDH15 EC10-MAD12 and dimeric PCDH15 EC9-MAD12 predict unrolling of MAD12 away from EC11 followed by MAD12 unfolding before EC repeats are perturbed (51, 87). As with PCDH15 EC8-10, we used an end-to-end colvar (N- and C-terminal residues) to obtain structural predictions and compared them to available crystal structures and stretched conformations obtained using SMD of fragments including MAD12 (51, 87).

All unsteered and steered predictions of PCDH15 EC9-MAD12 included three Ca^2+^ ions that were placed at their expected binding sites in the EC10-EC11 linker. In unsteered predictions, PCDH15 EC9-MAD12 exhibited a backbone RMSD of 2.5 ± 0.4 Å compared to the available structure (PDB: 6EET) (51) and showed the characteristic bent conformation of EC9-10 along with the folded MAD12. The predicted end-to-end distance was 117.1 ± 3.4 Å (Figure 3C, SFigure 5A). The sequence-overlap end-to-end distance was 106.3 Å for the crystallographic structure and 111.8 ± 3.4 Å in Boltz-2 predictions. With a harmonic restraint on the end-to-end distance of 130 Å, the distance was 125.4 ± 2.0 Å, but otherwise results were qualitatively similar to the unsteered predictions (Figure 3C, SFigure 5B). Applying a harmonic restraint of 160 Å to the end-to-end distance showed a distance of 151.3 ± 0.6 Å and straightening of the bent conformation for all predictions. In four of the five predictions, the post-MAD12 α helix at the C-terminal end showed some unfolding (Figure 3C, SFigure 5C). For a harmonic restraint of 210 Å, the end-to-end distance was 184.7 ± 3.6 Å. In all five predictions, the post-MAD12 α-helix at the C-terminal end showed unfolding, and the EC9-10 linker showed straightening (SFigure 5D). For a harmonic restraint of 260 Å, the end-to-end distances predicted were 206.7 ± 5.8 Å. Two predictions showed stretching of the EC9- 10 linker, while all predictions showed increased unfolding at the post-MAD12 α-helix. Two of these predictions also showed shredding at the C-terminal end (SFigure 5E). Altogether, we observed the general straightening of the EC9-10 linker which occurred along with unfolding of a terminal post-MAD12 α-helix (87). SMD simulations of PCDH15 EC10-MAD12 predicted the unfolding of this helix as well. However, our Bayesian-steered Boltz-2 predictions showed the unfolding of the post MAD12 helix and extraction of MAD12’s last β strand without unrolling of MAD12. SMD simulations instead predicted the unrolling of MAD12 before removal of that β strand (51, 87). This indicates that inertial or hydrodynamics effects arising in fast stretching SMD simulations with explicit water may not be captured by our approach when using a simple colvar with harmonic restraint.

### Predictions for the dimeric CDH23 and PCDH15 EC1-2 complex show unbinding with modest unfolding

The third inner-ear related system went beyond monomers and tested our approach on an antiparallel “handshake” dimer formed by the tips of CDH23 and PCDH15 (52). The CDH23 and PCDH15 EC1-2 hetero- dimeric complex (205 + 233 amino acids, *STable 1*) is essential for hearing and balance across vertebrates, it is under resting tension (88), and may break in response to large physiological stimuli such as loud sounds (51–58, 89, 90). The antiparallel dimer is formed by an overlap of the protein fragments in which EC1 from one monomer interacts with both EC1 and EC2 of the other. All-atom SMD simulations predict unbinding through sliding before unfolding, with a unique PCDH15 RGGPP loop acting as a hook preventing easy separation of the monomers from each other (52, 91). We used a colvar that involved C-terminal residues of each monomer to induce separation of the CDH23 and PCDH15 EC1-2 complex and compared our predictions to available crystal structures and unbinding trajectories obtained using SMD (52, 91).

In unsteered predictions, the antiparallel orientation of the CDH23 and PCDH15 EC1-2 dimer was correctly predicted in four out of five predictions, with these four predictions exhibiting a backbone RMSD of 5.1 ± 5.4 Å compared to the available structure (PDB: 4APX) (52). However, this estimate was heavily skewed by the fourth prediction that had a slightly different, yet still antiparallel, arrangement of monomers. The first three predictions had a backbone RMSD of 2.0 ± 0.4 Å compared to the deposited structure. The buried surface area (BSA) for these four same predictions was 1105.8 ± 205.6 Å^2^, in good agreement with the crystallographic value of 907 Å^2^. The fifth prediction failed to reproduce the expected antiparallel dimeric interface and placed the two monomers in parallel (Figure 3D SFigure 6A). In all these cases, Ca^2+^ ions were placed at expected binding sites. The predicted dimers’ end-to-end distance for the antiparallel complexes was 104.3 ± 1.9 Å. The sequence-overlap end-to-end distance was 103.9 Å for the experimental structure and 103.9 ± 1.8 Å in Boltz-2 predictions. Applying a harmonic restraint of 120 Å, led to a predicted end-to-end distance of 117.3 ± 0.8 Å. Most predictions showed slight separation at the interface with an antiparallel complex, but in one prediction, the interface was mostly lost (BSA of 227.2 Å), and the monomers were placed perpendicular to each other (Figure 3D SFigure 6B). The BSA for all complexes was 830.9 ± 350.5 Å^2^. Again, Ca^2+^ ions were placed at expected binding sites. For a harmonic restraint of 150 Å, the predicted end-to-end distance was 144.8 ± 1.4 Å. In all these predictions, Ca^2+^ ions were also placed at expected binding sites and the interface spanned only half the length of each monomer with EC1-EC1 contacts and a BSA of 274.8 ± 195.0 Å^2^. All five predictions also showed an interaction involving the RGGPP loop of PCDH15 (52). Three had a similar interface while the other two were each slightly different (Figure 3D SFigure 6C). This interaction was similar to what has been previously observed in SMD simulations (52). These results suggest that Bayesian-steered Boltz-2 may be able to predict intermediate states of the complex.

Predictions for the CDH23 and PCDH15 EC1-2 complex with a harmonic restraint set to 250 Å resulted in a predicted end-to-end distance of 227.8 ± 4.0 Å. All five predictions showed complete separation of the CDH23 and PCDH15 monomers with vanishing BSA. However, one prediction also exhibited unfolding in the CDH23 EC1-2 linker that kept the monomers in closer proximity compared to other predictions. A second prediction also exhibited unfolding in the CDH23 EC1-2 linker but no contacts between monomers (SFigure 6D). In this last case one Ca^2+^ ion was placed away from the expected binding residues. After increasing the harmonic restraint to 350 Å, the predicted end-to-end distance was 343.9 ± 1.4 Å (SFigure 6E). All of the predictions showed dramatic separation of the two monomers again with vanishing BSA, but both CDH23 and PCDH15 were intact with no unfolding. Unexpectedly, Ca^2+^ ions were either expelled from their binding sites or overlapped with multiple Ca^2+^ ions occupying the same site (Figure 3D, SFigure 6E). In SMD simulations, the monomers were observed to stay structurally intact but separated when subject to stretching forces (52). Our Bayesian-steered Boltz-2 predictions repeated this pattern with three out of five predictions at the harmonic restraint value of 𝑑 = 250 Å. Interestingly, for the larger separation value of 𝑑 = 350 Å, all monomers were intact for five out of five predictions. These results suggest that steered Boltz-2 predictions favor breaking the dimeric interface over compromising secondary structure of the monomers themselves.

### Predictions for the pentameric MscL show transitioning from a closed to an open-like state

Finally, to explore how Bayesian-steered Boltz-2 would predict the opening of multimeric ion channels, we applied colvar biases to the prokaryotic channel MscL and compared predictions to a crystal structure of its closed state (PDB: 2OAR) (92) and to experimentally constrained models of its open state (70–75). MscL is a pentamer that acts as an osmotic pressure sensor and safety release valve in bacteria where it is expected to undergo a large conformational change triggered by membrane tension alone (69–72). The open state has a conductance > 3 nS, which suggests that the open pore may have a radius between 14 Å and up to 20 Å (93). We used the sequence of MscL (151 x 5 = 755 amino acids, *STable 1*) and a colvar that involved the average distance of the center of mass of each valine 21 residue at the neck of the pore from their common center of mass. This distance has been previously used in enhanced sampling studies as a measure of the openness of the pore (73).

We ran five unsteered predictions using the same protocol as for previous systems. In these unsteered predictions, the pentameric MscL exhibited a backbone RMSD of 1.8 ± 0.2 Å compared to the available structure of the closed state (PDB:2OAR) (92). Its pore colvar was 5.1 ± 0.1 Å and its pore radius measured using HOLE (94) was found to be 1.0 ± 0.2 Å at its neck, consistent with a closed state (Figure 4, SFigure 7A, SFigure 8A, SMovie *1*). We additionally ran 500 unsteered predictions using Boltz-2’s original settings. These default predictions made by Boltz-2 only predict the closed state: across 500 samples the average distance of each valine 21 from their common center of mass was also 5.1 ± 0.1 Å. When steering predictions towards a possible transition state with a harmonic restraint of 10 Å, the exhibited pore colvar was 10.6 ± 0.3 Å and the neck radius was 6.6 ± 0.4 Å (Figure 4, SFigure 7B, SFigure 8B). Upon applying a harmonic restraint of 15 Å, the exhibited pore colvar was 15.1 ± 0.4 Å, and the neck radius was 10.3 ± 0.7 Å (Figure 4, SFigure 7C, SFigure 8C, SMovie *2*). Opening is achieved by helix tilting and expansion (helix tilt model), as opposed to expansion without tilting (barrel- stave model). We expect this MscL conformation to be representative of an open-like conformation. While a fully open pentameric structure of MscL is not available, several predictions based on electron paramagnetic resonance (EPR) (70), disulfide crosslinking (72), Förster resonance energy transfer (FRET) and MD simulations (71, 75) have been made. These experimentally constrained predictions consistently support open states like the one we predict with Bayesian-steered Boltz-2.

**Figure 4:**
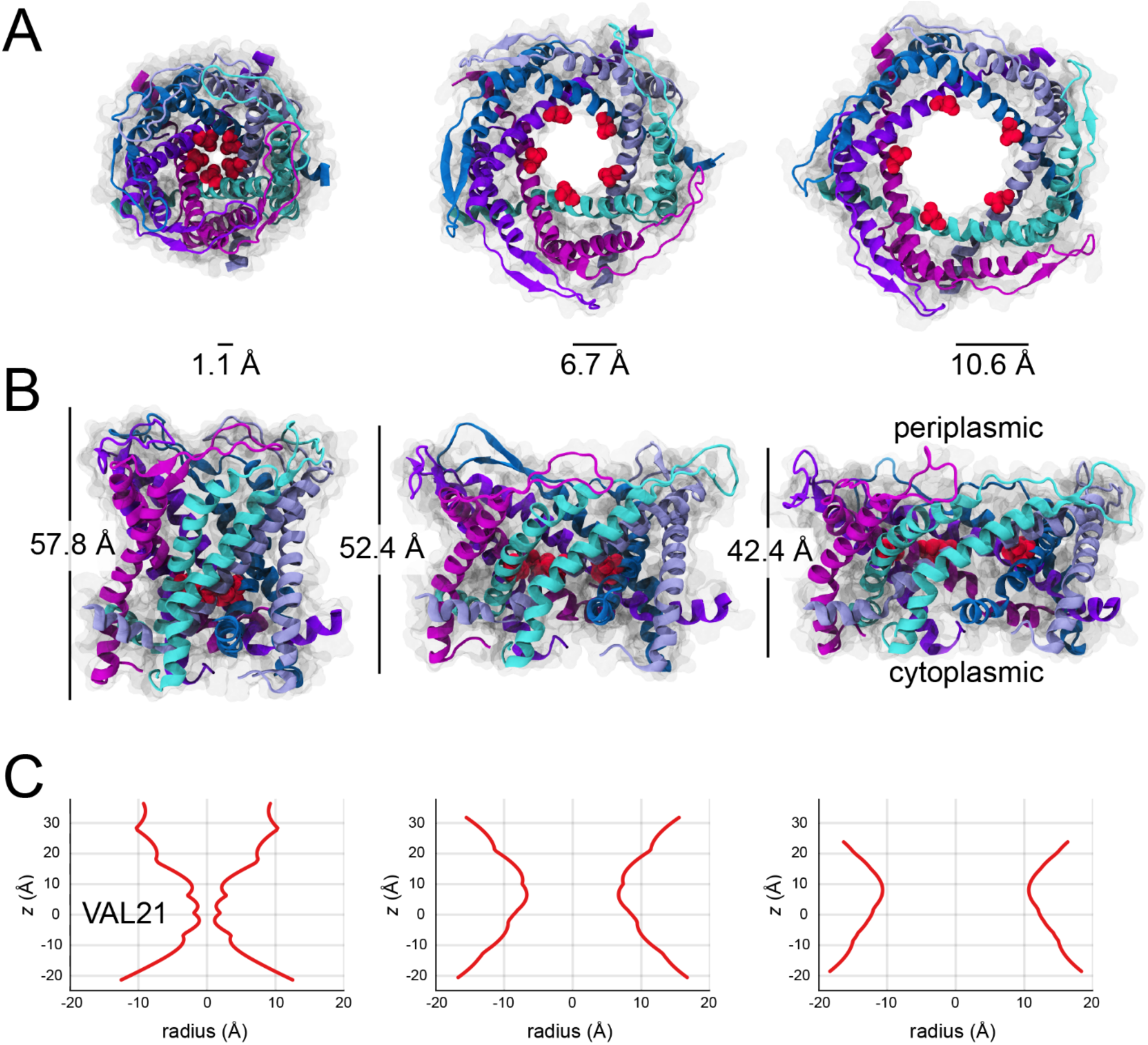
Bayesian-steered Boltz-2 predictions for the pentameric *Mycobacterium tuberculosis* MscL ion channel at varying values of the pore colvar. The colvar was defined as a vector whose five components were the distances of the centers of mass of each chain’s valine 21 residue from the common center of mass of all five residues. Each component of the colvar was unbiased (left column) and harmonically restrained at 10 Å (middle column) and 15 Å (right column). *A*. Top-down view of the pore opening. The radial value shown was computed using HOLE (94). Protein is shown in cartoon and transparent surface. Monomers are shown in cyan, ice blue, blue, violet, and magenta. Targeted valine 21 atoms at the neck of the pore are shown as red spheres. *B*. Side view of the pore opening. Cytoplasmic domain is omitted for visualization purposes (see SFigure 9). Figure panels in *A* and *B* are also shown in SFigure 7 and SFigure 8. *C*. Pore radii profiles for structure predictions in *A* and *B*. The *z*-coordinate has its origin at the center of mass of the valine 21 residues. The negative radial values are only intended as a reflection of the positive radial values on the right to facilitate visualization of the pore size. Pore radii were measured with HOLE (94) and are distinct from the pore colvar.

## Discussion

In this work, we have shown that Bayesian-steered twisted diffusion can produce alternative biomolecular conformations beyond those sampled natively by Boltz-2. Moreover, these conformations can be obtained using simple and intuitive Bayesian likelihoods (colvar biases). For each of our systems, we used harmonic restraints to prescribe target geometric distances. The resulting structure predictions were obtained at a fraction of the computational cost of traditional all-atom MD simulations. For all our colvars, at any fixed harmonic restraint the time to run our predictions ranged from only ∼ 10 min to ∼ 11.5 h on a 32 GB NVIDIA RTX-5000. To validate these predictions, we compared them to available experimental data and preexisting SMD simulations.

We found that Bayesian-steered Boltz-2 predicts many structural features and conformations expected from experimental data and SMD simulations. Our examples focused on mechanical biomolecules and ranged from the untwisting of a double-stranded DNA fragment to unfolding and stretching pathways for titin and PCDH15 fragments, separation of the CDH23 and PCDH15 dimeric handshake, and prediction of the open state for the pentameric MscL ion channel. In all cases we found structural models that recapitulated what was expected from experiments and SMD simulations. However, there were also some discrepancies between Boltz-2 and SMD predictions. In several cases with discrepancies, it is possible that stretching to larger end-to-end distances and increasing the number of Feynman-Kac particles (samples) would increase consensus with SMD, such as with titin and SMD showing the extraction of the G 𝛽 strand away from the A’ 𝛽 strand or SMD showing the unrolling of the MAD in PCDH15 EC9-MAD12. Alternatively, discrepancies may indicate a difference between what Boltz-2 has learned of protein biophysics and what is expected from MD force fields, which are themselves physical approximations often subject to undersampling issues. Unfortunately, traditional confidence metrics like the predicted local distance difference test (pLDDT) and the predicted aligned error (pAE) are difficult to interpret when generating predictions of stretched biomolecules that are by design supposed to differ from those used in training data sets for AlphaFold. Although Feynman-Kac particle weights rank predictions relative to one another, the lack for confidence metrics still leaves comparisons with difficult-to-obtain experimental data on stretched biomolecules and with SMD simulations as the only resources to validate the predictions.

All our Bayesian-steered Boltz-2 predictions were obtained using simple colvars that were only harmonically restrained during twisted diffusion. A strength of the Bayesian framework is that it can be iterated, and more sophisticated colvars accounting for additional physics can be used to steer Boltz-2 predictions increasing their compatibility with a user’s expectations. In this sense, one may view our approach as making the best Boltz-2 structure predictions that can be made given available experimental information. Moreover, the relative complexity of the colvar needed to produce predictions consistent with experiments or simulations might be taken as an informal measure of how applicable the physics learned by Boltz-2 is. It is remarkable that Bayesian steering with the simple colvars used here is able to produce complex predictions such as the open state conformations for MscL that are consistent with previously published models. Our Bayesian-steered Boltz-2 predictions, however, do not provide information on dynamics, forces, and energetics (41, 50, 95–101). Furthermore, although we have presented our Bayesian-steered predictions in order of increasing stretched distances, this direction may not correspond with a forward physical “time”. The predicted biomolecular conformations may also represent states on paths that go in opposite “time” directions over the defined colvar without differentiating between mechanical unfolding/folding, unbinding/binding, and opening/closing pathways. It has not escaped our attention that carrying out MD simulations on the Bayesian-steered Boltz-2 predicted structures can help resolve these shortcomings. Altogether, we find that twisted diffusion can predict structures that are consistent with experiments, reproduce predictions from SMD simulations, and do this using simple and intuitive steering constraints while also being significantly cheaper in compute time compared to all-atom MD.

To steer Boltz-2 towards predicting rare conformations, we reframed this quest as a sampling problem drawing from a conditioned diffusion distribution. We used a likelihood set by the user to define a posterior predictive distribution with the Boltz-2 foundation model as a prior, but sampling from this posterior presents its own technical obstacles. To overcome them, we used twisted diffusion (32) and Feynman-Kac steering (33) approaches, which are based on sequential Monte Carlo (30, 35). Like most sampling methods, there are several important hyperparameters that can affect performance. First, there are the resampling strategy and number of Feynman-Kac particles used during steering. We implemented stratified resampling instead of the default multinomial resampling because of its improved, reduced variance properties (30, 35, 102). Because of the high dimensional nature of distributions defined over atomic coordinates, an expected limitation is that particle weight degeneracy may occur. It is therefore practical to think of the number of particles in relation to a compute versus prediction quality tradeoff. One may use as many particles as needed until the predictions are of a sufficient quality whether that be assessed by the variance of the predictions as the number of particles is increased or by expert knowledge. As few as two particles can yield improvements in sample quality (32, 33).

Two additional hyperparameters that can affect steered Boltz-2 prediction performance are the tempering coefficients used on the particle weights, α in equation (*17*), and the steering gradients, γ in equation *(1S)*. In principle, their values do not matter as the number of particles grows to infinity (30, 34, 35), but for a finite number of particles their choices can affect convergence rates and prediction quality. We expect that optimal choices will depend on how well the true Bayes steering term is approximated by the proposal term (equation (*22*)), which is likely application specific. In this work, we identified a common set of tempering coefficients α and γ that were used across all our predictions. The last hyperparameter we highlight is the number of diffusion steps. We have empirically found that increasing the number of diffusion steps reduces the shredding that was more prevalent in predictions using stronger constraints. This trend is consistent with general numerical solvers requiring smaller integration steps to resolve dynamics under large steering gradients or flow fields since diffusion models are regarded as discretizations of stochastic differential equations (28). Investigations into the sensitivity of predictions on hyperparameters will be valuable, but our examples show that reasonable hyperparameter settings already resulted in physically accurate predictions across a range of systems.

To further illustrate the effectiveness of twisted diffusion steering using Boltz-2, we address its computational feasibility. We have found that restrictions set by GPU memory, whether due to system size and the number of residues or the number of Feynman-Kac particles, were the most practical limitation of our method. However, the memory demands placed by the number of particles can be mitigated by reducing the number predicted in parallel. Also, we modified Boltz-2 to not only compute gradients over denoised coordinates when steering but to additionally compute gradients of the likelihood through the denoiser. This requires pytorch to build an automatic differentiation computational graph during steering which also increases the use of GPU memory. In our applications, we did not find this memory consumption to be limiting comparatively. With all these technical qualifications in mind, we have found that our approach is practical for systems with up to at least ∼ 3,000 residues on high end GPUs.

Interestingly, we found that modifying Boltz-2 to compute guidance gradients through the denoiser improved its ability to make significant global rearrangements such as those required to predict the open-like state of MscL. We hypothesize this is due to the denoiser containing significant information, learned during training, about how residues are correlated with one another. This may permit relatively simple likelihoods that only explicitly involve a few residues to propagate implications across many more residues. On the other hand, computing guidance gradients only on denoised coordinates means there is little to no correlation information between residues except what is explicitly provided in the likelihood function. This hypothesis warrants future investigation.

As we shape and refine biomolecular structure prediction tools, we expect that these will shape our quest to understand dynamical structure-function relationships. Characterizing the structural and biophysical properties of biomolecules under tension has been historically challenging. Initial single-molecule force-spectroscopy experiments provided forces and extensions with little information on structural conformations at atomistic resolution. SMD simulations have provided predictions at high spatial and time resolution, albeit with limitations related to force-field accuracy and timescales. Some of these predictions have been experimentally verified thus providing insights into the micromechanics of biomolecules (66). In parallel, a few ad-hoc experimental approaches have been developed and used to capture stretched states of mechanical proteins and mechanosensitive membrane proteins in ways that are amenable for high-resolution structure determination (103–110). Yet these approaches remain difficult to implement and generalize. Here we provide an alternate *in*-*silico* tool for fast and accurate prediction of geometrically “stretched” biomolecular structures. We expect that this tool will facilitate the generation of testable hypothesis about the mechanics of biomolecules and the exploration and design of stretch-dependent binding partners for functional tests and therapeutics. Broadly, Bayesian-steered Boltz-2 should facilitate the exploration of structural conformations along any colvar, including those defining the functional states of adhesion proteins, transporters, ion channels, ribosomes, and any biomolecular complex undergoing conformational changes during assembly or function.

A major strength of the method shown here is its flexibility and adaptability for most any application. Since the diffusion model is steered by an arbitrary Bayesian likelihood, virtually any statistical relationship or collective variable bias between the atomic coordinates and user-supplied constraints or experimental data can be imposed. This approach is natural and part of a larger emerging paradigm shift whereby foundation models are used as Bayesian priors because of their state-of-the-art status in sample quality (111). This paradigm shift is driving rapidly emerging efforts on developing both enhanced conformational sampling and improved biomolecular structure modeling using generative-AI (112–120). Our work complements these efforts both through the applications we have demonstrated and by its implementation inside of Boltz-2, an AlphaFold3-like foundation model that can predict multimeric structures comprised of proteins, DNA, RNA, and ligands. We have presented examples of the natural and broadly applicable case of conditioning by harmonic restraints, but like the enhanced sampling methods that continue to be developed for MD simulations, we anticipate opportunities for innovating and refining application-specific steering methods. We expect that these approaches will facilitate investigations exploring novel biomolecular conformations that are rare under the default configuration of Boltz-2 and related foundation models.

## Methods

Boltz-2’s generative structure module with physics-based corrections is a weighted Markov process that is conditioned on the sequencing information:

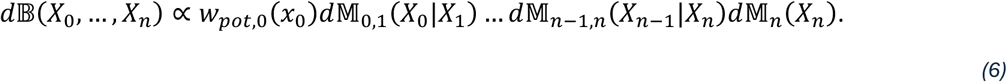

The generative process is represented by the probability distribution on diffusion trajectories 𝑑𝔹. For technical reasons, here the variables 𝑋_𝑘_represent an extended state vector that includes the Gaussian noised atomic coordinates at diffusion steps 𝑡_𝑘_and *t̂*_𝑘_(see **Appendix I: Methods**). We let 𝑥_0_ stand for noiseless coordinates at diffusion step 0. The weight function 𝑤_𝑝𝑜𝑡,0_(𝑥_0_) was defined by Boltz-2 (7) using its logarithm and imposes energetic penalties on non-physical structural poses. The Markov kernels 𝑑𝕄_𝑘−1,𝑘_(𝑋_𝑘−1_|𝑋_𝑘_), which are probability distributions for the variable 𝑋_𝑘−1_ conditioned on the value of 𝑋_𝑘_, represent Boltz-2’s reverse EDM step from the noisy structure 𝑋_𝑘_ to the slightly less noisy 𝑋_𝑘−1_. Unlike a standard EDM, these kernels also apply random rotations and translations to the structures (see **Appendix I: Methods**).

We obtain a posterior distribution by conditioning equation *(C)* on an auxiliary variable 𝜉, multiplying by the usual Bayesian likelihood 𝑝(𝜉|𝑥_0_):

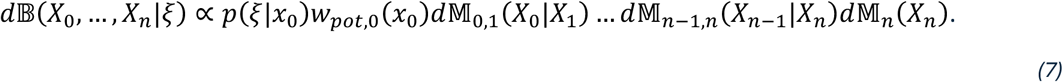

The right hand side of equation (*7*) is the unnormalized, resulting posterior which defines a probability distribution on diffusion trajectories, 𝑑𝔹(𝑋_0_, …, 𝑋_𝑛_|ξ), that is conditioned on a fixed value of ξ selected by the user. We choose functions 𝐺 and proposal kernels 𝑑ℚ so that the posterior distribution is written as a Feynman-Kac model (34, 35):

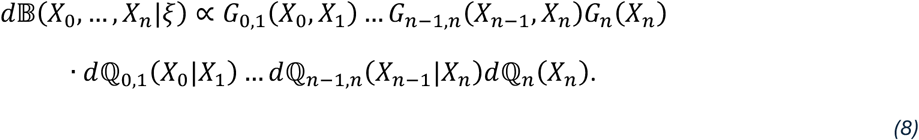

Our choices for *G* and 𝑑ℚ are based on a twisted diffusion sampler (32) (see **Appendix I: Methods**) and belong to the broader class of Feynman-Kac steered diffusion models (33). A weighted sample approximation to d𝔹(X_0_, …, X_n_|ξ) using 𝓅 Feynman Kac particles is produced through the particle filter algorithm (35):

Boltz-2 (7) implemented a particle filter algorithm with multinomial resampling instead of stratified resampling. This was done for a fundamentally different Feynman-Kac model that, as one example, did not condition on an arbitrary Bayesian likelihood. We have implemented Algorithm 1 for equation (*8*) inside of the August 25^th^, 2025, nightly version of Boltz-2 (git commit #0c228c). This required custom implementation of the functions 𝐺 and proposal kernels 𝑑ℚ prescribed by twisted diffusion (32). We provide further details, including changes to several of Boltz-2’s default hyperparameter values, in the supplement (see **Appendix I: Methods**).

### Algorithm 1: Pseudocode for the standard SMC sampler associated to the Feynman-Kac model in equation (8).

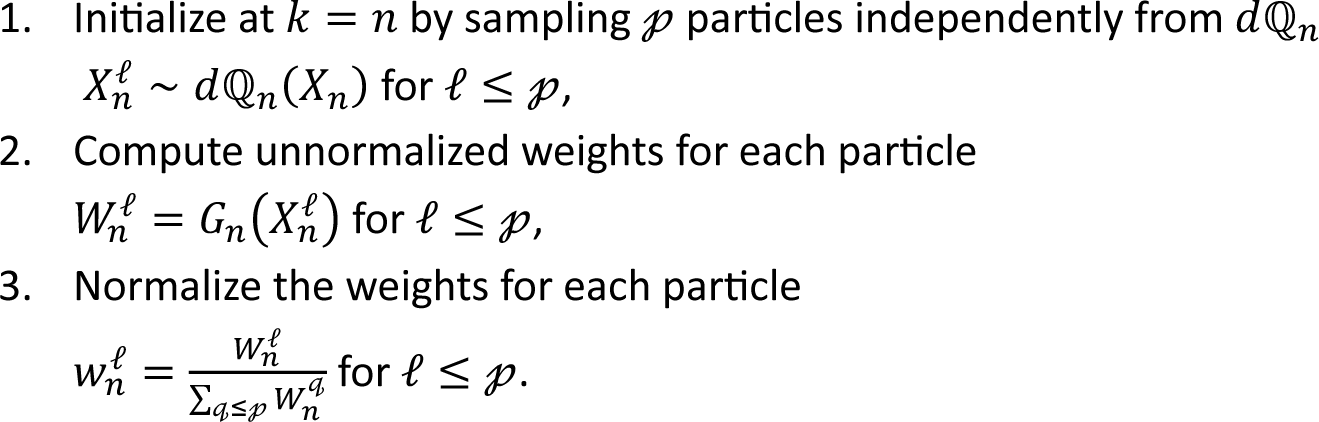
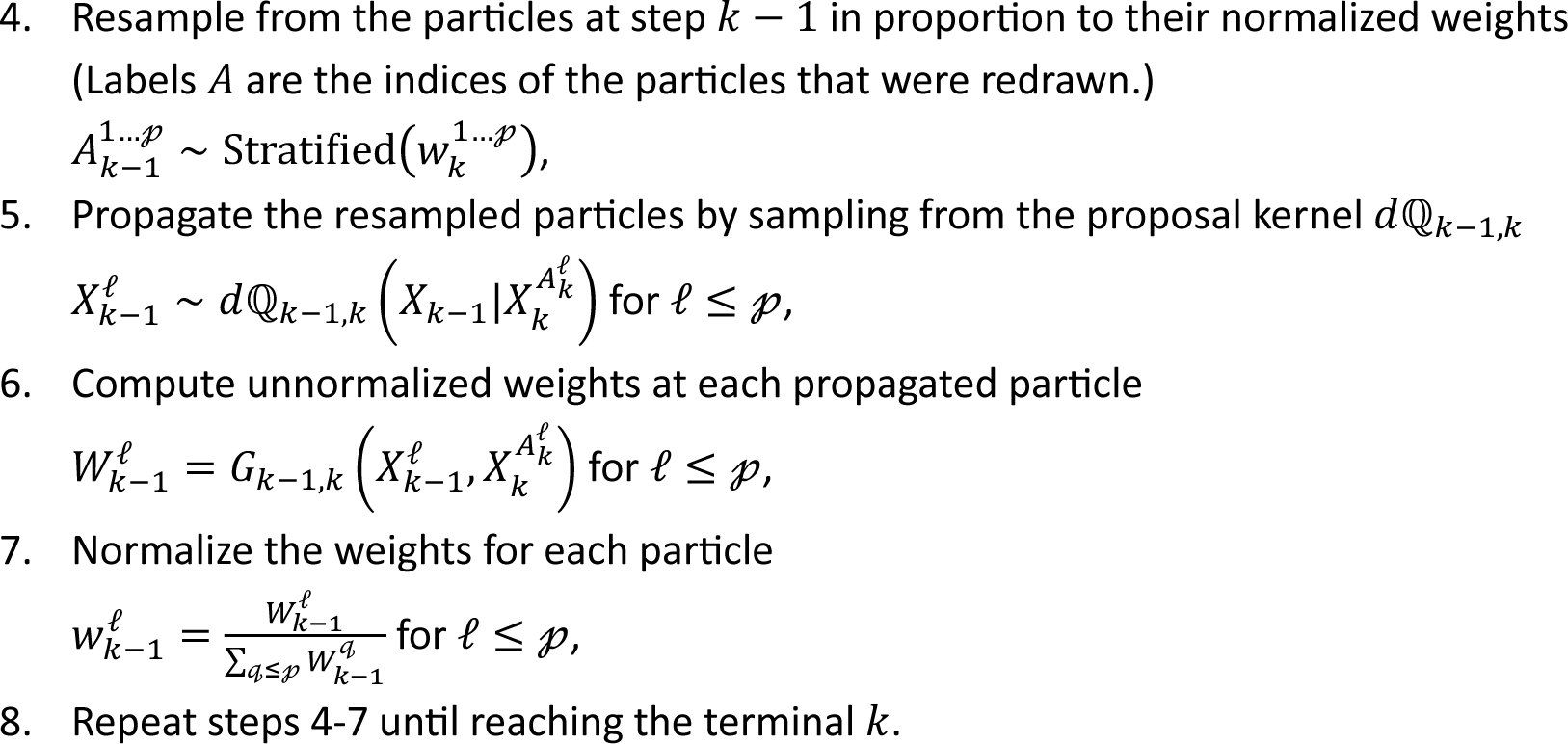

For the systems we analyzed, we used Bayesian-steered Boltz-2 and sequences to make five independent predictions (𝑛 = 5) each using 100 Feynman-Kac particles. We reported the highest probability particle for prediction output. For MscL we made an additional 500 predictions using Boltz-2’s original settings to demonstrate that, by itself, Boltz-2 exclusively predicts the closed state. We stress that the biomolecules were only represented with sequence information. No templates or additional structural information were used. No additional training or re-training of Boltz-2 was required. For all steered predictions, we used a normally distributed Bayesian likelihood whose mean was a colvar we designed as a function of the atomic coordinates (*e.g.,* distances between atoms). Equivalently, we harmonically restrained the value of this colvar. We sought to restrain distances between the centers of mass of atom selections, but rather than restrain those distances directly, we passed them through sigmoid functions and obtained a unitless colvar 𝜉 whose value was 1 when the target distances were attained. We then equivalently restrained 𝜉 to be 1. Restraining the target distances directly led to larger gradients which could destabilize the generative diffusion without mitigation. Unless otherwise stated, the standard deviation used to enforce a harmonic restraint was 0.01 and the number of diffusions steps taken was 200.

The DNA sequence used in our predictions was a 12-bp fragment based on the initial portion of λ-DNA, that was stretched in early single-molecule force spectroscopy experiments (40). Protein sequences used in our predictions (*STable 1*) were obtained from structures in the protein data bank: titin (PDB: 1TIT), PCDH15 EC8-10 (PDB: 4XHZ), PCDH15 EC9-MAD12 (PDB:6EET), CDH23 EC1-2 and PCDH15 EC1-2 (PDB: 4APX), and MscL (PDB:2OAR). For all but DNA, MscL, and the CDH23 and PCDH15 EC1-2 complex, when restraints were applied, we restrained the centers of mass of the N- and C-terminal residues to be at a target distance. Similarly, for the CDH23 and PCDH15 EC1-2 complex, restraints were applied to the center of mass of the C-terminal residues. For titin, these distances were 60 Å − 75 Å in steps of 5 Å. For PCDH15 EC8-10, PCDH15 EC9-MAD12, and CDH23 EC1-2 and PCDH15 EC1-2 we increased the target distance in increments of 20 Å, 50 Å, 100 Å, and 150 Å relative to its native Boltz-2 value. Stretching predictions for PCDH15 EC8-10 at 160 Å, 210 Å, and 260 Å used 400 diffusion steps. In the case of no Ca^2+^, the stretching at 130 Å also used 400 diffusion steps. For the dimeric interface between CDH23 and PCDH15, we increased the target distance in increments of 20 Å, 50 Å, 150 Å, and 250 Å relative to Boltz- 2’s native value. These predictions all used a harmonic restraint standard deviation of 0.0025. All predictions with Ca^2+^ included three ions per canonical linker region and one extra ion for the tip of CDH23 EC1 (121) and the truncated tip of PCDH15 EC9. For the harmonic restraint at the largest distance, we also used 2,000 diffusion steps. For MscL, we restrained the centers of mass of each of the valine residues at the pore neck (residue 21) to be at a distance of 10 Å and 15 Å from their common center of mass. For the latter prediction, we used 400 diffusion steps. Pore radii, distinct from the pore colvar, were computed using HOLE (94). BSA was computed using VMD (122), which was also used to prepare all molecular images. The lion images (Figure 1A) are wholly for conceptual illustration and were not generated by a full twisted diffusion pipeline. Instead, ChatGPT (123) was used to create the two terminal lion images. We then forward noised the images but displayed them in the denoising direction, since in the limit that the forward and reverse diffusion processes perfectly match, this corresponds to conditioning the reverse trajectories on their terminal value.

## Code availability

Our Bayesian-steered Boltz-2 is available for download on github along with documentation (https://github.com/Sotomayorlab-UChicago/Public-BayesianSteeredBoltz2). Users are expected to adjust the `compute_variable` and `compute_function` methods within the `BayesianPotential` subclass defined in the `colvars.py` file. To aid users in defining their own colvars, we have also created a Jupyter notebook called `Boltz2AtomIds.ipynb`. The first section of this notebook reconstructs Boltz- 2’s internal representation of residues by atom index from the protein sequence, e.g., FASTA or YAML file. The relevant atom indices from this notebook should be passed to `compute_variable` to define the desired colvar. The second section exports a `connected_atom_index.pt` tensor that automates the inclusion of the backbone restraint potential along the phosphodiester bond for DNA and the C-N peptide bond for proteins.

## Author contributions

C.K. conceived Boltz modifications and M.S. guided work on biomolecular case studies. C. K. wrote all code and ran all simulations. C. K. and M.S. wrote the manuscript.

## Acknowledgments

This work was supported in part by the University of Chicago and the National Institutes of Health through the National Institute on Deafness and Other Communication Disorders (R01-DC015271 to M.S.). Simulations were performed using local and ACCESS resources at the National Center for Supercomputing Applications (Delta supercomputer, grant ACCESS MCB140226 to M.S.). This research used the Delta advanced computing and data resource which is supported by the National Science Foundation (award OAC 2005572) and the State of Illinois. Delta is a joint effort of the University of Illinois Urbana-Champaign and its National Center for Supercomputing Applications. We gratefully acknowledge discussions with Dr. Benoit Roux, Dr. Harper E. Smith, Haosheng Wen, and members of the Roux and Sotomayor laboratories.

## Notes

### Competing Interest Statement

The authors have declared no competing interest.

