## Supplement for "Bayesian-Steered Structure Prediction of Mechanical Biomolecules Using Twisted Diffusion"

SMovie 1: **Unsteered denoising diffusion trajectory for MscL (closed state)**. Standard Boltz-2 prediction for the pentameric MscL. Protein is primarily shown in licorice representation with the valine 21 atoms at the eventual neck of the pore shown as red spheres. Movie shows the secondary structure gradually forming. Final frames show the predicted structure in cartoon and transparent surface. Monomers are shown in cyan, ice blue, blue, violet, and magenta.

SMovie 2: **Bayesian-steered denoising diffusion trajectory for MscL (open-like state).** Bayesian-steered Boltz-2 prediction for the pentameric MscL. The colvar involving valine 21 residues was harmonically restrained at 15 Å. Protein is shown as in SMovie 1.

### Appendix I: Methods

**Conditioning Boltz's diffusion distribution by a Bayesian likelihood**

The diffusion module implemented in AlphaFold3 and Boltz-2 is based on the elucidated diffusion model (EDM) (76) but has been heuristically modified (3, 6, 7) to promote rotational and translational invariance in the structure predictions. At an index $k$, the diffusion module samples

$$\mathscr{r}_k \text{ – the frame of reference rotation and translation variables at index } k,$$

$$x_{t_k} \text{ – the noised atomic coordinates at step } t_k,$$

$$x_{\hat{t}_k} \text{ – the noised atomic coordinates at step } \hat{t}_k. \tag{9}$$

We collect these variables into a single state vector at index $k$:

$$X_k = \left(x_{\hat{t}_k}, x_{t_k}, \mathscr{r}_k\right). \tag{10}$$

The EDM is a generative model, based on the variance exploding formulation of diffusion (26, 28, 76), which alternates between denoising steps from $x_{\hat{t}_k}$ to $x_{t_{k-1}}$ along the deterministic reverse ODE flow and noising steps from $x_{t_{k-1}}$ to $x_{\hat{t}_{k-1}}$ along the stochastic forward Brownian motion (76). The step indices are ordered as $t_{k-1} \leq \hat{t}_{k-1} \leq \hat{t}_k$ . For the EDM, the forward noising schedule is taken as $\sigma(t) = t$ where $\sigma(t)$ is the standard deviation of the normally distributed $p(x_t|x_0)$ at step $t$. This generative process is Markovian, and we notate Boltz-2's transition kernels as

$$d\mathbb{M}_n(X_n) = p_{\hat{n}}\left(x_{\hat{t}_n}|x_{t_n}, \mathscr{r}_n\right) dx_{\hat{t}_n} p_n\left(x_{t_n}\right) dx_{t_n} p_R(\mathscr{r}_n) d\mathscr{r}_n,$$

$$d\mathbb{M}_{k-1,k}(X_{k-1}|X_k) = p_{\widehat{k-1}}\left(x_{\hat{t}_{k-1}}|x_{t_{k-1}}, \mathscr{r}_{k-1}\right) dx_{\hat{t}_{k-1}} d\delta_{\Psi_{k-1,k}\left(x_{\hat{t}_k}\right)}\left(x_{t_{k-1}}\right) p_R(\mathscr{r}_{k-1}) d\mathscr{r}_{k-1}. \tag{11}$$

The density $p_R(\mathscr{r})$ over the frame of reference variables corresponds to how Boltz-2 samples a random rotation and translation vector. The density $p_n\left(x_{t_n}\right)$ is a normal distribution centered at the origin. Its standard deviation is $t_n$ in correspondence with the EDM's noise schedule. In Boltz-2, the deterministic function $\Psi_{k-1,k}\left(x_{\hat{t}_k}\right)$ resembles an Euler step approximation along the reverse ODE flow starting at step $\hat{t}_k$ and terminating at step $t_{k-1}$ so that the conditional distribution for $x_{t_{k-1}}$, notated as $d\delta_{\Psi_{k-1,k}\left(x_{\hat{t}_k}\right)}\left(x_{t_{k-1}}\right)$, is a Dirac mass:

$$\Psi_{k-1,k}\left(x_{\hat{t}_k}, D_{\hat{t}_k}\right) = \mathrm{WRA}\left(x_{\hat{t}_k}, D_{\hat{t}_k}\right) + \eta\left(\frac{t_{k-1} - \hat{t}_k}{\hat{t}_k}\right)\left(\mathrm{WRA}\left(x_{\hat{t}_k}, D_{\hat{t}_k}\right) - D_{\hat{t}_k}\right),$$

$$\Psi_{k-1,k}\left(x_{\hat{t}_k}\right) = \Psi_{k-1,k}\left(x_{\hat{t}_k}, D\left(x_{\hat{t}_k}, \hat{t}_k\right)\right). \tag{12}$$

The function $D\left(x_{\hat{t}_k}, \hat{t}_k\right)$ is the denoised prediction for $x_{\hat{t}_k}$ at step $\hat{t}_k$. The function $\mathrm{WRA}\left(x_{\hat{t}_k}, D_{\hat{t}_k}\right)$ is a weighted-rigid-alignment between its arguments, and its use is again a heuristic modification of the EDM made by AlphaFold3 and Boltz-2 (3, 6, 7) to promote rotational and translational invariance. The step-scale parameter η is another heuristic modification by AlphaFold3 and Boltz-2 (3, 6, 7). If WRA were the identity on $x_{\hat{t}_k}$ and η were 1, then equation *(12)* would be an Euler step along the reverse ODE flow (76). The density $p_{\widehat{k-1}}\left(x_{\hat{t}_{k-1}} | x_{t_{k-1}}, \mathscr{r}_{k-1}\right)$ is the normal distribution for forward noising which in accordance with the EDM noising schedule has a variance given by $\hat{t}_{k-1}^2 - t_{k-1}^2$, which Boltz-2 heuristically scaled by a factor 1.006. Its mean is given by

$$\mu_{\widehat{k-1}}\left(x_{t_{k-1}}, \mathscr{r}_{k-1}\right) = r_{k-1}\left(x_{t_{k-1}} - \overline{x_{t_{k-1}}}\right) + tr_{k-1}, \tag{13}$$

where $r_{k-1}$ is the random rotation and $tr_{k-1}$ is the random translation vector that are both represented by $\mathscr{r}_{k-1}$. The expression $\overline{x_{t_{k-1}}}$ stands for the center of mass of the atom coordinates. We will also sometimes shorthand the right-hand side of equation *(13)* as $\mathscr{r}_{k-1} \cdot x_{t_{k-1}}$.

Boltz-2 further implemented physics-based potentials (7) for mitigating steric clashes and other hallucination phenomena to which diffusion models are prone (3, 124). These physics-based potentials are encoded as a Boltzmann weight factor $w_{pot,0}(x_0)$ applied to the overall joint distribution so that the standard Boltz-2 generative model with physics corrections is given by

$$d\mathbb{B}(X_0, \dots, X_n) \propto w_{pot,0}(x_0) d\mathbb{M}_{0,1}(X_0|X_1) \dots d\mathbb{M}_{n-1,n}(X_{n-1}|X_n) d\mathbb{M}_n(X_n). \tag{14}$$

Note equation *(14)* here coincides with equation *(6)* in the main text. We remark that this diffusion distribution is internally conditioned by Boltz-2 on the evolutionary sequence information coming from the multiple sequence alignments (MSA) and trunk module (7), but for our purposes that is a static parameter, and we suppress that dependence in our notation.

We condition equation *(14)* on an auxiliary variable $\xi$ by multiplying by the usual Bayesian likelihood $p(\xi|x_0)$

$$d\mathbb{B}(X_0, \dots, X_n|\xi) \propto p(\xi|x_0) w_{pot,0}(x_0) d\mathbb{M}_{0,1}(X_0|X_1) \dots d\mathbb{M}_{n-1,n}(X_{n-1}|X_n) d\mathbb{M}_n(X_n). \tag{15}$$

Equation *(15)* also coincides with equation *(7)* in the main text. The likelihood function $p(\xi|x_0)$ can be any arbitrary probability density and is determined by the desired application.

**The twisted diffusion sampler**

Sequential Monte Carlo (SMC) algorithms (30, 35) are randomized sampling schemes for computing normalized and unnormalized integrals of the type

$$\int \dots \int f(x_n) H_{n,n-1}(x_n, x_{n-1}) \dots H_{1,0}(x_1, x_0) H_0(x_0) d\mathbb{P}_{n,n-1}(x_n|x_{n-1}) \dots d\mathbb{P}_{1,0}(x_1|x_0) d\mathbb{P}_0(x_0). \tag{16}$$

The nonnegative functions $H_{k,k-1}$ are called potentials, and the associated measures are called Feynman-Kac models (34). Note that the negative logarithm of an SMC potential is more analogous to the physics concept of a potential energy. We will use the term *potential* to refer to the SMC definition and *potential energy* to refer to its negative logarithm. Our conditioned diffusion model equation *(15)* is of this form up to trivially relabeling the indices in descending order and realizing that the predictive distribution for atomic coordinate structures is the marginal at step $t = 0$. There is freedom to pick the potentials and Markov transition kernels so long as their product returns equation *(15)*. Going forward we will notate our potentials and transition kernels in descending order as $G_{k-1,k}$ and $d\mathbb{Q}_{k-1,k}$. We will also refer to the latter kernels as proposal kernels.

We follow recent work on designing Feynman-Kac models for conditioning diffusion models (32, 33). Our choices of potentials and transition kernels are based on those of a twisted diffusion sampler (32) but adjusted for the heuristic modifications made for rotational and translational invariance and Boltz-2's physics-based potential. Additionally, we include tempering parameters. We first consider the weight factor in equation *(15)* and rewrite it in the telescoping, *i.e.,* twisted, form

$$p(\xi|x_0)w_{pot,0}(x_0) =$$
$$\left(\prod_{k=n}^{1} \frac{p\left(\xi|D\left(x_{\hat{t}_{k-1}},\hat{t}_{k-1}\right)\right)^{\alpha_{k-1}}}{p\left(\xi|D\left(x_{\hat{t}_k},\hat{t}_k\right)\right)^{\alpha_k}} \frac{w_{pot,k-1}\left(D\left(x_{\hat{t}_{k-1}},\hat{t}_{k-1}\right)\right)}{w_{pot,k}\left(D\left(x_{\hat{t}_k},\hat{t}_k\right)\right)}\right) p\left(\xi|D\left(x_{\hat{t}_n},\hat{t}_n\right)\right)^{\alpha_n} w_{pot,n}\left(D\left(x_{\hat{t}_n},\hat{t}_n\right)\right).$$

*(17)*

Note this right-hand side telescoping pattern more precisely returns the conditioning weight factor $p\left(\xi|D\left(x_{\hat{t}_0},\hat{t}_0\right)\right) w_{pot,0}\left(D\left(x_{\hat{t}_0},\hat{t}_0\right)\right) \approx p(\xi|x_0)w_{pot,0}(x_0)$. We expect no practical difference between the use of either of these weight factors in equation *(15)* since in our numerical implementation these terminal diffusion steps Gaussian-noise the protein structure with a standard deviation on the order of $0.002$ Å.

The functions $w_{pot,k}$ were already implemented by Boltz-2, and we keep their definition. The powers $\alpha$ are tempering hyperparameters which may be chosen by the user to fine-tune SMC performance. The only restriction is that $\alpha_0 = 1$.

For the proposal kernels, we set

$$d\mathbb{Q}_n(X_n) = p_{\hat{n}}\left(x_{\hat{t}_n}|x_{t_n},r_n\right)dx_{\hat{t}_n}p_n\left(x_{t_n}\right)dx_{t_n}p_R(r_n)dr_n,$$

$$\mathrm{d}\mathbb{Q}_{k-1,k}(\mathrm{X}_{k-1}|\mathrm{X}_k) = \mathrm{q}_{\widehat{k-1}}\left(x_{\hat{t}_{k-1}}|x_{\hat{t}_k},r_{k-1}\right)\mathrm{d}x_{\hat{t}_{k-1}}\mathrm{d}\delta_{\Psi_{k-1,k}\left(x_{\hat{t}_k}\right)}\left(x_{t_{k-1}}\right)\mathrm{p_R}(r_{k-1})\mathrm{d}r_{k-1}.$$

*(18)*

The only difference between equations *(11)* and *(18)* are the conditional kernels on $dx_{\hat{t}_{k-1}}$ which now depend on $x_{\hat{t}_k}$ instead of $x_{t_{k-1}}$. In particular, $d\mathbb{M}_{k-1,k}(X_{k-1}|X_k) = \frac{p_{\widehat{k-1}}\left(x_{\hat{t}_{k-1}}|x_{t_{k-1}},r_{k-1}\right)}{q_{\widehat{k-1}}\left(\mathrm{x}_{\hat{t}_{k-1}}|\mathrm{x}_{\hat{t}_k},r_{k-1}\right)} d\mathbb{Q}_{k-1,k}(X_{k-1}|X_k)$. We define the proposal kernel in terms of the original as

$$q_{\hat{t}_{k-1}}\left(x_{\hat{t}_{k-1}}|x_{\hat{t}_k},r_{k-1}\right) = p_{\hat{t}_{k-1}}\left(x_{\hat{t}_{k-1}}|\Phi_{k-1,k}\left(x_{\hat{t}_k}\right),r_{k-1}\right),$$

$$\Phi_{k-1,k}(x_{\hat{t}_k}) = \Psi_{k-1,k}\left(x_{\hat{t}_k}, D(x_{\hat{t}_k}, \hat{t}_k) - \nabla_{x_{\hat{t}_k}} E_k(x_{\hat{t}_k})\right),$$

$$E_k(x_{\hat{t}_k}) = -\ln w_{grd,k}\left(D(x_{\hat{t}_k}, \hat{t}_k)\right) - \gamma_k \hat{t}_k^2 \ \ln p\left(\xi | D(x_{\hat{t}_k}, \hat{t}_k)\right). \tag{19}$$

In keeping with classifier guidance techniques (28, 31, 32), the denoised prediction within the $\Psi_{k-1,k}$ function has been perturbed by the gradient of a potential energy function defined by the Bayesian likelihood as well as Boltz-2's physics potentials. In particular, the log-likelihood plays the role of the classifier. While Boltz-2 was implemented to take gradients of $E_k$ with respect to the denoised coordinates $\nabla_{D_{\hat{t}_k}}$, we take gradients with respect to noisy coordinates $\nabla_{x_{\hat{t}_k}}$ and therefore include the additional step of computing gradients through the denoiser function as is done in other inference time steering applications (32). Like $w_{pot,k}$, the functions $w_{grd,k}$ were also already implemented in Boltz-2, and we keep those definitions. Although not the same, the functions $w_{pot,k}$ and $w_{grd,k}$ are similar. In both cases and for all $k$, their negative logarithm is spanned by a common set of basis functions such as energetic penalty terms on bond distances and dihedral angles. Their negative logarithms only differ in their scalar coefficients across those basis functions, with those scalar coefficients again playing the role of tempering hyperparameters. Similarly, the $\gamma$ coefficients are hyperparameters for tempering the likelihood gradient and to fine-tune the SMC performance.

The $\hat{t}_k^2$ weight coefficient on the log-likelihood term in equation *(19)* is derived from how the denoiser transforms under conditioning due to its relationship with the score function, $\nabla \ln p\,(x_t)$. For the variance exploding SDE, the denoiser and score satisfy (76)

$$D(x_t, t) = x_t + \sigma^2(t)\nabla_{\mathrm{x_t}} \ln p\,(x_t). \tag{20}$$

Under conditioning, the score transforms according to Bayes' formula: $\nabla_{\mathrm{x_t}} \ln p\,(x_t|\xi) = \nabla_{\mathrm{x_t}} \ln p\,(x_t) + \nabla_{\mathrm{x_t}} \ln p\,(\xi|x_t)$. which when combined with equation *(20)* and the EDM's noise schedule $\sigma(t) = t$, implies

$$D(x_t, t|\xi) = D(x_t, t) + t^2\nabla_{\mathrm{x_t}} \ln p\,(\xi|x_t). \tag{21}$$

By abuse of notation, we have let $D(x_t, t|\xi)$ stand for the denoiser of the conditioned diffusion process even though the denoiser itself is not a probability density. Finally, $p(\xi|x_t)$ stands for the conditional density of $\xi$ given the noisy atomic coordinates at step $t$ which is intractable. We therefore use the classifier guidance-like approximation (32)

$$\nabla_{\mathrm{x_t}} \ln p(\xi|x_t) \approx \nabla_{\mathrm{x_t}} \ln p\left(\xi|D(x_t, t)\right) \tag{22}$$

where the latter likelihood is now the user prescribed likelihood for $\xi$ given the denoised atomic coordinates $D(x_t, t)$ at step 0. In principle, SMC can correct the approximation error here in the asymptotic large sample limit. In this work, we did not put the $\hat{t}_k^2$ weight factor on the $\log w_{grd,k}$ term in equation *(19)* so as to make minimal changes with Boltz-2's standard use of $w_{grd,k}$.

Combining equations *(15)*, *(17)*, and *(18)* we arrive at our twisted diffusion, Feynman-Kac model for our target distribution:

$$\int \dots \int f(x_0) d\mathbb{B}(X_0, \dots, X_n | \xi) \propto$$
$$\int \dots \int f(x_0) G_{0,1}(X_0, X_1) \dots G_{n-1,n}(X_{n-1}, X_n) G_n(X_n) d\mathbb{Q}_{0,1}(X_0 | X_1) \dots d\mathbb{Q}_{n-1,n}(X_{n-1} | X_n) d\mathbb{Q}_n(X_n),$$

$$G_n(X_n) = p\left(\xi | D\left(x_{\hat{t}_n}, \hat{t}_n\right)\right)^{\alpha_n} w_{pot,n}\left(D\left(x_{\hat{t}_n}, \hat{t}_n\right)\right),$$

$$G_{k-1,k}(X_{k-1}, X_k) = \left(\frac{p\left(\xi | D\left(x_{\hat{t}_{k-1}}, \hat{t}_{k-1}\right)\right)^{\alpha_{k-1}}}{p\left(\xi | D\left(x_{\hat{t}_k}, \hat{t}_k\right)\right)^{\alpha_k}} \frac{w_{pot,k-1}\left(D\left(x_{\hat{t}_{k-1}}, \hat{t}_{k-1}\right)\right)}{w_{pot,k}\left(D\left(x_{\hat{t}_k}, \hat{t}_k\right)\right)}\right) \frac{p_{\widehat{k-1}}\left(x_{\hat{t}_{k-1}} | x_{t_{k-1}}, r_{k-1}\right)}{q_{\widehat{k-1}}\left(x_{\hat{t}_{k-1}} | x_{\hat{t}_k}, r_{k-1}\right)}. \quad (23)$$

**Numerical implementation**

We have implemented *Algorithm 1* for equation *(23)* inside of the August 25th, 2025, nightly version of Boltz-2 (git commit #0c228c). Going forward, our discussion of Boltz-2 and any changes we made refer to that version.

The most important changes we made to Boltz-2 were to compute the likelihood terms in equations *(23)* and *(19)* and incorporate them into Boltz-2's implementation of *Algorithm 1*. For this purpose, we defined a `BayesianPotential` subclass of Boltz-2's `Potential` class in our `colvars.py` file. Its `compute_variable` method returns any function of the atomic coordinates that is needed to compute the likelihood function. For example, if distances between atomic coordinates were being harmonically restrained, this method would compute those distances. We refer to these intermediate functions of the atomic coordinates as collective variables (colvars). The `compute_function` method returns the negative log of the desired Bayesian-likelihood as a function of the outputs of `compute_variable`. Computing the negative log-likelihood is internally consistent with Boltz-2's use of the `compute_function` method to return an energy for its physics-based potentials and Boltz-2's implementation of *Algorithm 1*.

The `BayesianPotential` subclass also implements a `compute` method that is internally used by Boltz-2 to compute the potential energy as a function of the supplied atomic coordinates. It essentially wraps sequential calls of `compute_variable` and `compute_function`. Similarly, this subclass implements a `compute_gradient` method that is internally used to compute the energy gradient with respect to the supplied atomic coordinates. Boltz-2's physics potentials were simple enough that their gradients could be computed analytically. To allow for arbitrarily defined likelihoods, we compute these gradients using pytorch's automatic differentiation routines (125).

Inside the `colvars.py` file, we have also implemented a `fkpf_options` function. This function returns a `parameters` dictionary which contains entries for the tempering schedules used for $\alpha$ in equation *(17)* and γ in equation *(19)*. For these tempering schedules, we have defined a `PowerSchedule` subclass of Boltz-2's `ParameterSchedule` class. The latter class is used by Boltz-2 to define tempering parameters that vary with steps during SMC as shown in the `compute_parameters` method of Boltz-2's `Potential` class. Our subclass of tempering schedules is modeled after $(1-x)^p$ on the interval $[0,1]$ but appropriately rescaled to a custom interval $[a,b]$. When `PowerSchedule` is declared with `flag_guidance_weight=True`, the tempering schedule includes the $t^2$ weight factor in equation *(19)*. Our `PowerSchedule` subclass is also defined in `colvars.py`.

We have integrated this customized `BayesianPotential` into Boltz-2's workflow by augmenting Boltz-2's `get_potentials` function. When our new `--use_colvars` flag is passed at the command line call of Boltz-2, the `get_potentials` function will now include the `BayesianPotential` class in its list of potentials for running SMC. Additionally, the output of `fkpf_options` is passed as parameters in the construction of the `BayesianPotential`.

In addition to defining the `BayesianPotential` class, it was necessary to modify several other internal features of Boltz-2's default SMC sampler. One of the most significant changes was to implement the $\nabla_{x_{\widehat{t_k}}}$ operator in equation *(19)*. In Boltz-2's `diffusionv2.py`, energy gradients are computed with respect to atomic coordinates after the denoiser has been applied. To pull these back by chain-rule to energy gradients with respect to noisy atomic coordinates, we used pytorch's automatic differentiation (125) to build a computational graph through the denoiser and `torch.autograd.grad` to compute a vector-Jacobian product with the energy gradient on denoised atomic coordinates. We also implemented stratified resampling instead of Boltz-2's default multinomial resampling (35, 102). Another notable change concerned the SMC potentials in equations *(23)*, *(24)*, *(25)*. Boltz-2 omitted the difference of WRA terms in equation *(25)* while we have reincluded it. (**Appendix II: Incorporating Difference of Weighted-Rigid Alignments term in the Boltz-2 computation of SMC potentials**). Finally, we explicitly populated Boltz-2's `ContactPotential` with bonds for every contiguous 3' oxygen and 5' phosphate atoms along DNA and for every C-N pair along the backbone within each polypeptide chain.

**Prediction protocols**

The predictions presented in this work, both steered and unsteered, all used a common set of hyperparameters except for minor changes (declared in **Methods**). We comment here on the common hyperparameters which differed from Boltz-2's default values. Boltz-2 defined the relationship between $t_k$ and $\hat{t}_k$ as $\hat{t}_k = (1+\gamma)t_k$. Instead of Bolt-2's maximum value of $\gamma = 0.8$, we took the value $0.05$. This choice was motivated by the fact that the reverse step in equation *(12)* is approximately an Euler step along the reverse ODE flow. The smaller increment was expected to lead to better approximations over smaller steps which mitigated large perturbative gradients introduced by the Bayesian likelihood. At the other extreme, instead of a minimum value of $\gamma = 0$, we took the value $10^{-6}$. This ensured that the proposal densities for $x_{\hat{t}_k}$ in equation *(23)* were continuous while $\gamma$ was essentially $0$ numerically. Boltz-2 also took a default value of $1.5$ for their $step\_scale$ parameter which is η in equation *(12)*. We took a value of $1.0$ so that equation *(12)* more closely resembled an Euler step along the reverse flow. Boltz-2's parameter $guidance\_interval$ permits a multi-step gradient descent procedure on the denoised prediction to find a perturbation towards a more optimal energy minimum. However, we have reset the number of gradient descent steps to $guidance\_interval = 1$. While higher values of $guidance\_interval$ are natural from an optimization point of view, from the standpoint of conditioning diffusion processes, a single gradient step may be more natural owing to equation *(21)*. We also set $fk\_resampling\_interval = 1$ since otherwise particle weights should be accumulated between resampling steps which Boltz-2 does not do by default. We finally changed the `ContactPotential`'s buffer parameter from $2$ Å to $1.75$ Å which penalized contact bonds to fall below $1.75$ Å.

Our predictions were derived from normally distributed Bayesian likelihoods $p(\xi|x_0)$ whose mean functions were colvars of the atomic coordinates $x_0$. In analogy with MD simulations (36-39), this amounted to placing harmonic restraints on the associated colvars. Our colvars were derived from centers of mass taken over appropriate atom selections, computing distances between those centers of mass, and applying sigmoidal functions to these distances. These sigmoidal functions were modeled after $3x^2 - 2x^3$ truncated to the interval $[0,1]$. Together, we used these operations to define colvars that were $1$ when our centers of mass were at the desired target distances and decayed to $0$ as these centers of mass were not at the target distance and harmonically restrained these colvars to be $1$ by corresponding normally distributed Bayesian likelihoods. We found the use of sigmoidal functions led to smaller perturbative gradients in equation *(21)*, while using the distances directly led to larger gradients that could be more destructive to the reverse diffusion unless mitigated by the computational expense of increased diffusion steps.

### Appendix II: Incorporating Difference of Weighted-Rigid Alignments term in the Boltz-2 computation of SMC potentials

Boltz-2's `diffusionv2.py` file omits a term in its computation of the SMC potentials equation *(23)* and the log-ratio of proposal densities. We reproduce the derivations for the missing term and have reincorporated it into our `diffusionv2.py` file.

We examine the ratio of target and proposal densities in equation *(23)*. Taking the log, we obtain

$$ln\left(\frac{p_{\widehat{k-1}}\left(x_{\hat{t}_{k-1}}|x_{t_{k-1}},r_{k-1}\right)}{q_{\widehat{k-1}}\left(x_{\hat{t}_{k-1}}|x_{\hat{t}_k},r_{k-1}\right)}\right)=\frac{-1}{2\left(\hat{t}_{k-1}^2-t_{k-1}^2\right)}\left(\left(x_{\hat{t}_{k-1}}-r_{k-1}\cdot\Psi_{k-1,k}\left(x_{\hat{t}_k},D\left(x_{\hat{t}_k},\hat{t}_k\right)\right)\right)^2-\left(x_{\hat{t}_{k-1}}-r_{k-1}\cdot\Psi_{k-1,k}\left(x_{\hat{t}_k},D\left(x_{\hat{t}_k},\hat{t}_k\right)-\nabla_{x_{\hat{t}_k}}E\left(x_{\hat{t}_k}\right)\right)\right)^2\right).$$

If we set $\epsilon = x_{\hat{t}_{k-1}}-r_{k-1}\cdot\Psi_{k-1,k}\left(x_{\hat{t}_k},D\left(x_{\hat{t}_k},\hat{t}_k\right)-\nabla_{x_{\hat{t}_k}}E\left(x_{\hat{t}_k}\right)\right)$, then this becomes

$$ln\left(\frac{p_{\widehat{k-1}}\left(x_{\hat{t}_{k-1}}|x_{t_{k-1}},r_{k-1}\right)}{q_{\widehat{k-1}}\left(x_{\hat{t}_{k-1}}|x_{\hat{t}_k},r_{k-1}\right)}\right)=\frac{-1}{2\left(\hat{t}_{k-1}^2-t_{k-1}^2\right)}\left(\left(\epsilon+r_{k-1}\cdot\Psi_{k-1,k}\left(x_{\hat{t}_k},D\left(x_{\hat{t}_k},\hat{t}_k\right)-\nabla_{x_{\hat{t}_k}}E\left(x_{\hat{t}_k}\right)\right)-r_{k-1}\cdot\Psi_{k-1,k}\left(x_{\hat{t}_k},D\left(x_{\hat{t}_k},\hat{t}_k\right)\right)\right)^2-\epsilon^2\right). \tag{24}$$

Multiplication by $r_{k-1}$ represents an affine transformation since centering by the center of mass and rotating are linear operations. Thus, $r_{k-1}\cdot u-r_{k-1}\cdot v=\mathcal{L}_{k-1}\cdot u-\mathcal{L}_{k-1}\cdot v$ for the linear operator, $\mathcal{L}_{k-1}$, composed of centering and rotating. Now we rewrite equation *(12)* as a linear combination of its terms:

$$\Psi_{k-1,k}\left(x_{\hat{t}_k},D_{\hat{t}_k}\right)=\left(1+\eta\left(\frac{t_{k-1}-\hat{t}_k}{\hat{t}_k}\right)\right)WRA\left(x_{\hat{t}_k},D_{\hat{t}_k}\right)-\eta\left(\frac{t_{k-1}-\hat{t}_k}{\hat{t}_k}\right)D_{\hat{t}_k}.$$

Combining these with equation *(24)* , distributing $\mathcal{L}_{k-1}$ across the linear combination above, and then regrouping terms as differences of the same type under $\mathcal{L}_{k-1}$, we obtain

$$r_{k-1}\cdot\Psi_{k-1,k}\left(x_{\hat{t}_k},D\left(x_{\hat{t}_k},\hat{t}_k\right)-\nabla_{x_{\hat{t}_k}}E\left(x_{\hat{t}_k}\right)\right)-r_{k-1}\cdot\Psi_{k-1,k}\left(x_{\hat{t}_k},D\left(x_{\hat{t}_k},\hat{t}_k\right)\right)$$

$$=\left(1+\eta\left(\frac{t_{k-1}-\hat{t}_k}{\hat{t}_k}\right)\right)\mathcal{L}_{k-1}\cdot\left(WRA\left(x_{\hat{t}_k},D\left(x_{\hat{t}_k},\hat{t}_k\right)-\nabla_{x_{\hat{t}_k}}E\left(x_{\hat{t}_k}\right)\right)-WRA\left(x_{\hat{t}_k},D\left(x_{\hat{t}_k},\hat{t}_k\right)\right)\right)-\left(\eta\left(\frac{t_{k-1}-\hat{t}_k}{\hat{t}_k}\right)\right)\mathcal{L}_{k-1}\cdot\left(-\nabla_{x_{\hat{t}_k}}E\left(x_{\hat{t}_k}\right)\right).$$

(25)

The first term containing the difference of weighted-rigid alignments was the omitted term referenced at the beginning of this section and is now included in our code. Boltz-2 also scales the variance by an additional factor 1.006 that we have omitted in the derivation here but include in the numerical implementation.

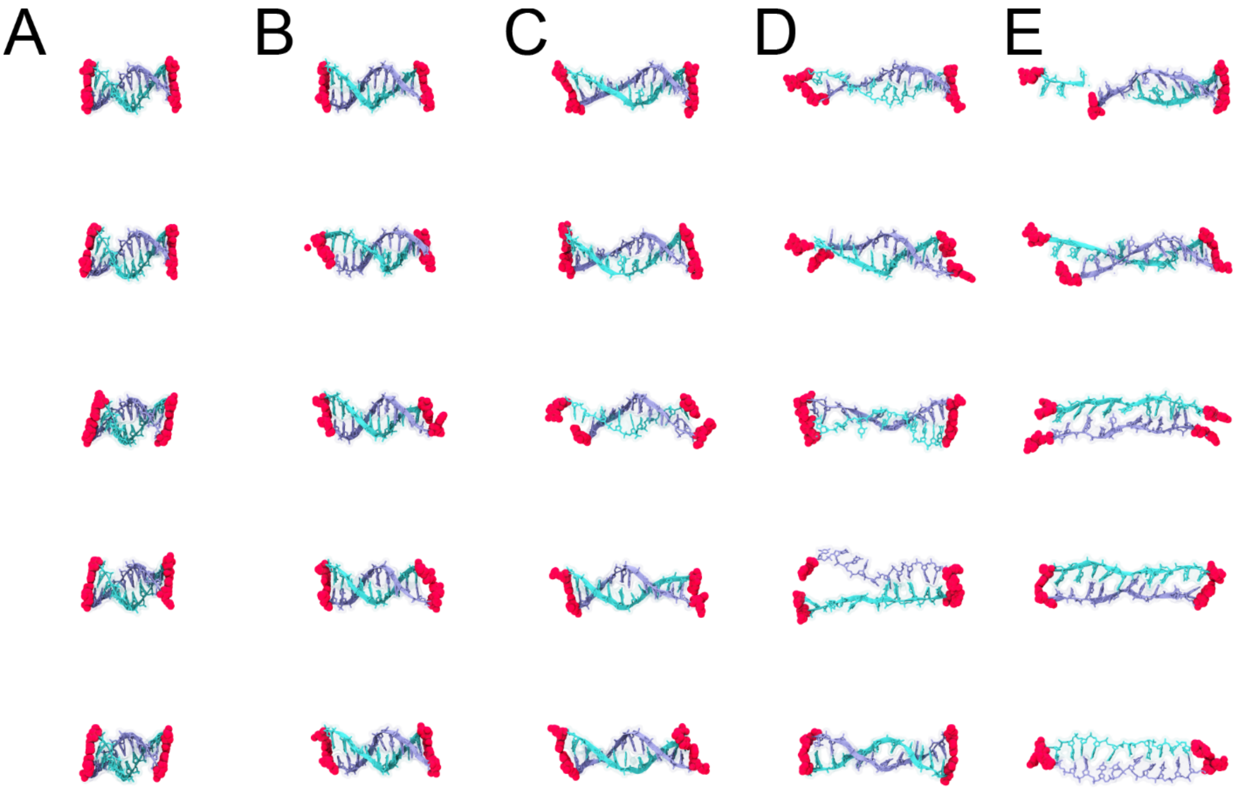

**SFigure 1: Bayesian-steered Boltz-2 predictions for DNA (GGGCGGCGACCT) at varying end-to-end distances.** The colvar was defined as the distance between centers of mass of each pair of terminal bases. Each prediction was generated using its own collection of 100 Feynman-Kac particles and selected by taking the resulting highest probability particle. *A*. Unsteered Boltz-2 predictions whose colvars natively take an approximate value of 35 Å. *B-E*. Bayesian-steered Boltz-2 predictions with the colvar harmonically restrained at 45 Å, 55 Å, 65 Å, and 75 Å, respectively. Shredding is observed in *E* (top). DNA shown as in Figure 2A. Some panels are also shown in Figure 2.

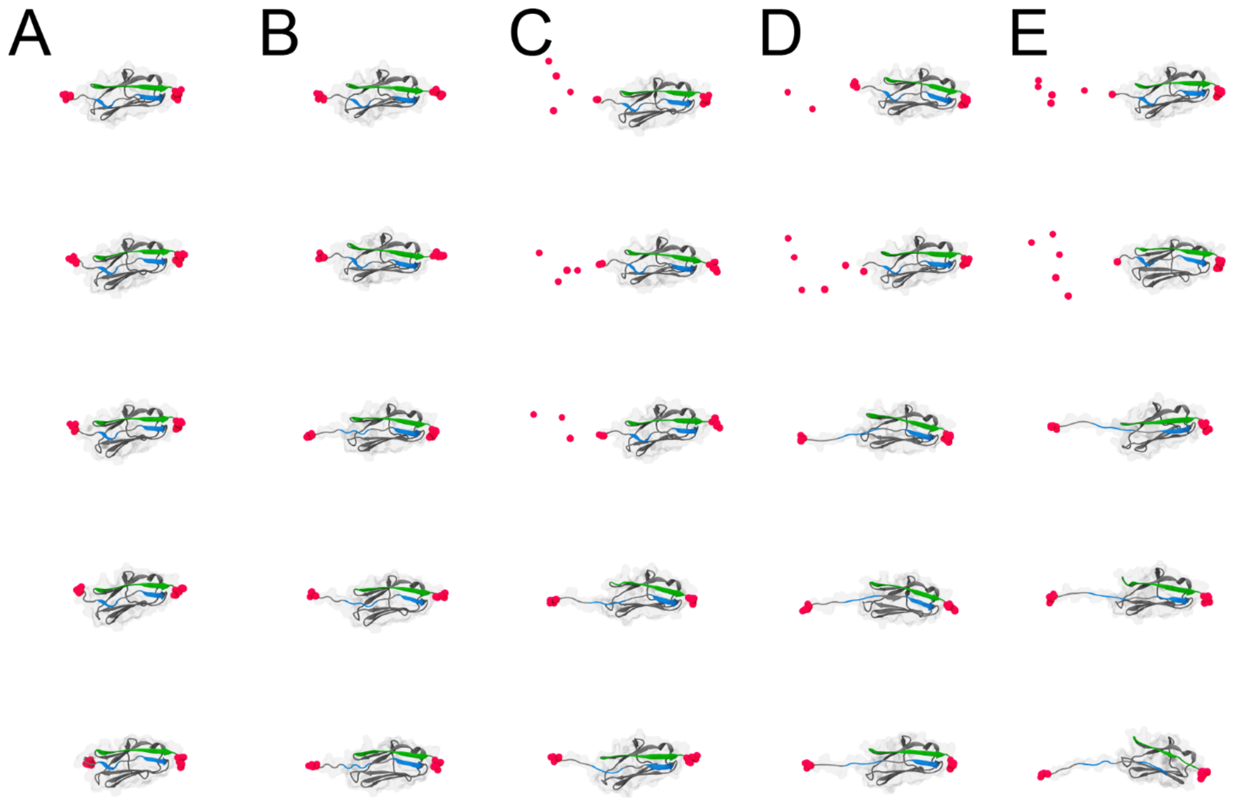

**SFigure 2: Bayesian-steered Boltz-2 predictions for titin I91 at varying end-to-end distances.** The colvar was defined as the distance between centers of mass of N- and C-terminal residues. Each prediction was generated using its own collection of 100 Feynman-Kac particles and selected by taking the resulting highest probability particle. *A*. Unsteered Boltz-2 predictions whose colvars natively take an approximate value of 55 Å. *B-E*. Bayesian-steered Boltz-2 prediction with the colvar harmonically restrained at 60 Å, 65 Å, 70 Å, and 75 Å, respectively. Shredding is observed at N-terminal residue for some predictions. Protein shown as in Figure 2B. Some panels are also shown in Figure 2.

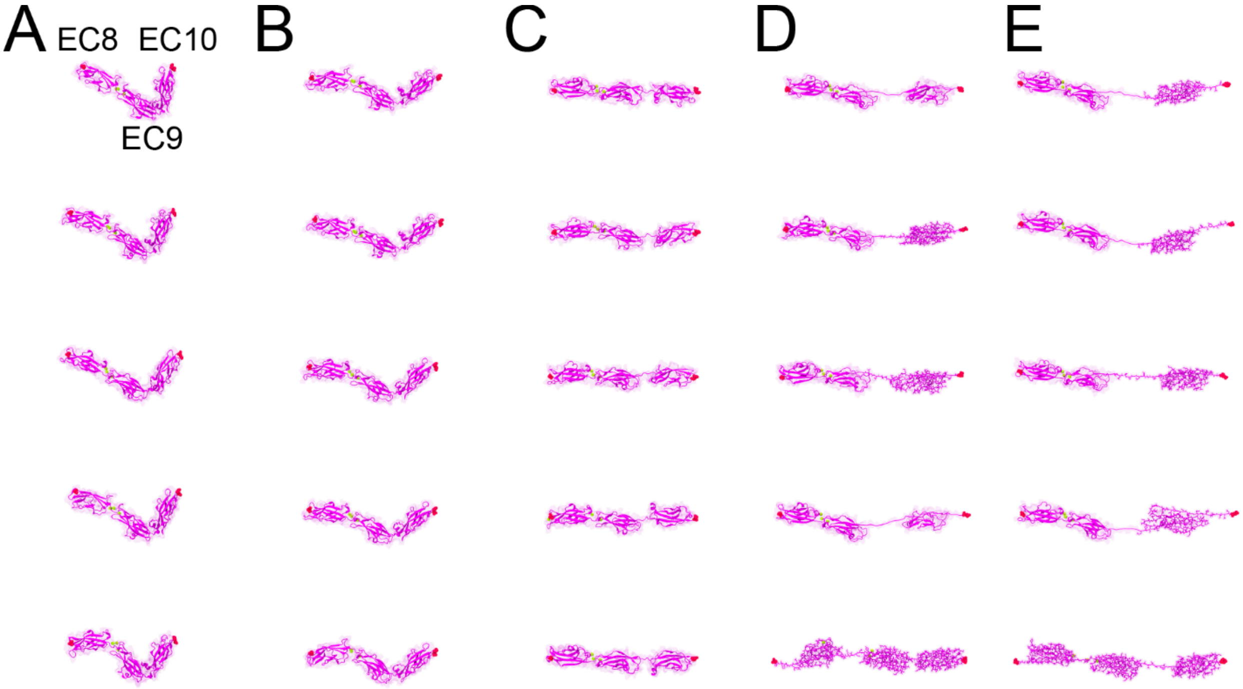

**SFigure 3: Bayesian-steered Boltz-2 predictions for PCDH15 EC8-10 with $Ca^{2+}$ at varying end-to-end distances.** The colvar was defined as the distance between centers of mass of N- and C-terminal residues. Each prediction was generated using its own collection of 100 Feynman-Kac particles and selected by taking the resulting highest probability particle. *A*. Unsteered Boltz-2 predictions whose colvars natively take an approximate value of 110 Å. *B*-*E*. Bayesian-steered Bolzt-2 predictions with the colvar harmonically restrained at 130 Å, 160 Å, 210 Å, and 260 Å, respectively. Protein shown as in Figure 3. Some panels are also shown in Figure 3.

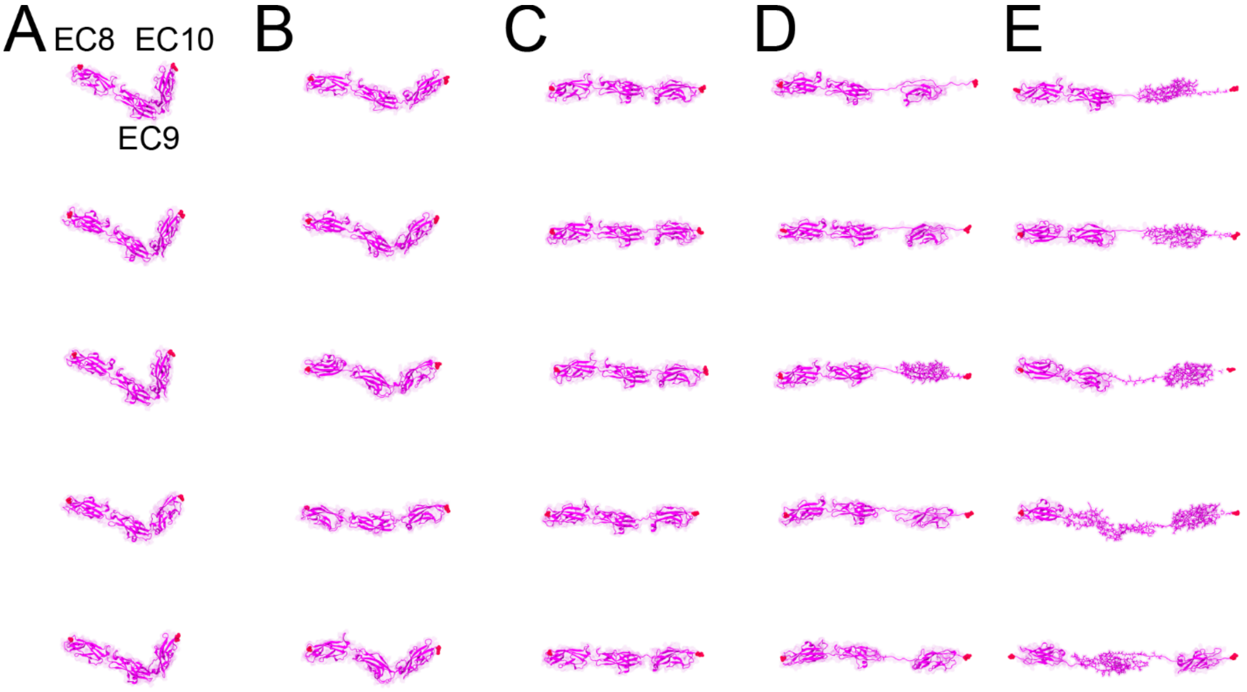

**SFigure 4: Bayesian-steered Boltz-2 predictions for PCDH15 EC8-10 without $Ca^{2+}$ at varying end-to-end distances.** The colvar was defined as the distance between centers of mass of N- and C-terminal residues. Each prediction was generated using its own collection of 100 Feynman-Kac particles and selected by taking the resulting highest probability particle. *A*. Unsteered Boltz-2 predictions whose colvars natively take an approximate value of 110 Å. *B-E*. Bayesian-steered Boltz-2 predictions with the colvar harmonically restrained at 130 Å, 160 Å, 210 Å, and 260 Å, respectively. Protein shown as in Figure 3. Some panels are also shown in Figure 3.

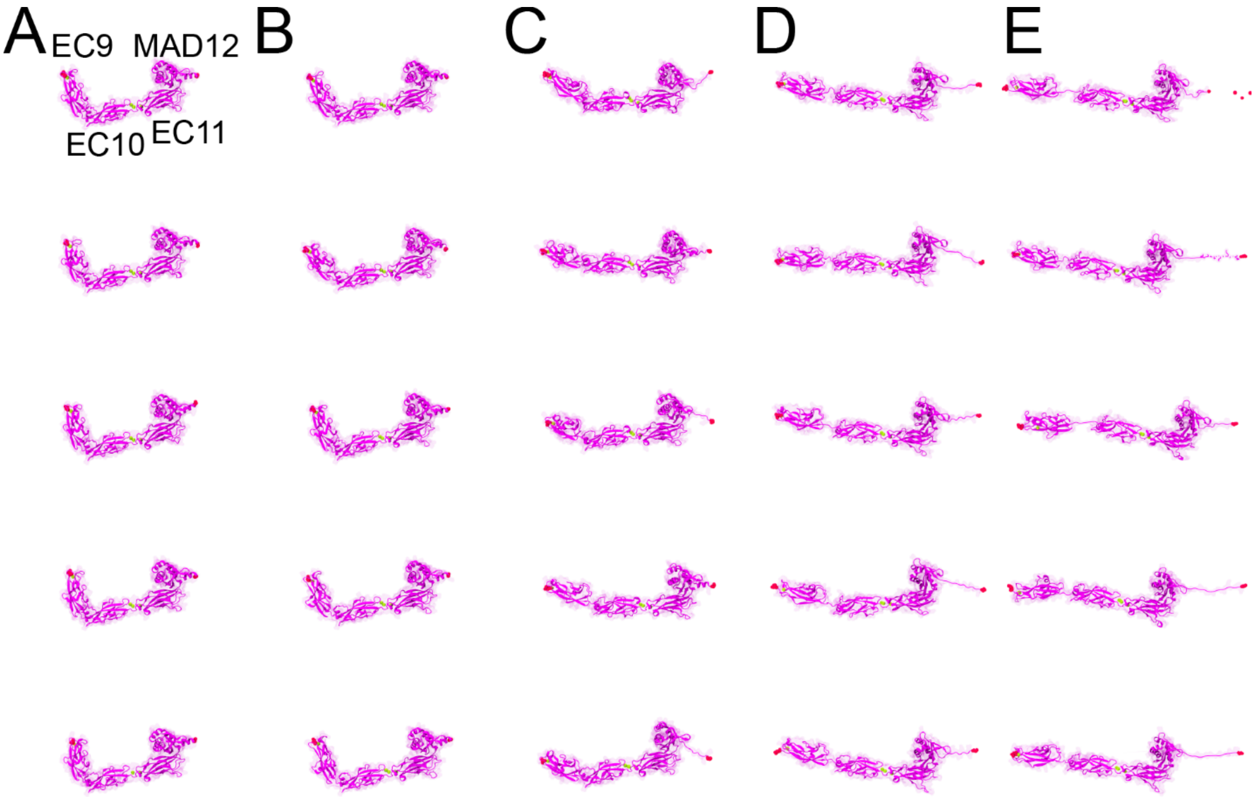

**SFigure 5: Bayesian-steered Boltz-2 predictions for PCDH15 EC9-MAD12 at varying end-to-end distances.** The colvar was defined as the distance between centers of mass of N- and C-terminal residues. Each prediction was generated using its own collection of 100 Feynman-Kac particles and selected by taking the resulting highest probability particle. *A*. Unsteered Boltz-2 predictions whose colvars natively take an approximate value of 110 Å. *B-E*. Bayesian-steered Boltz-2 predictions with the colvar harmonically restrained at 130 Å, 160 Å, 210 Å, and 260 Å, respectively. Shredding is observed at C-terminal residue for some predictions. Protein shown as in Figure 3. Some panels are also shown in Figure 3.

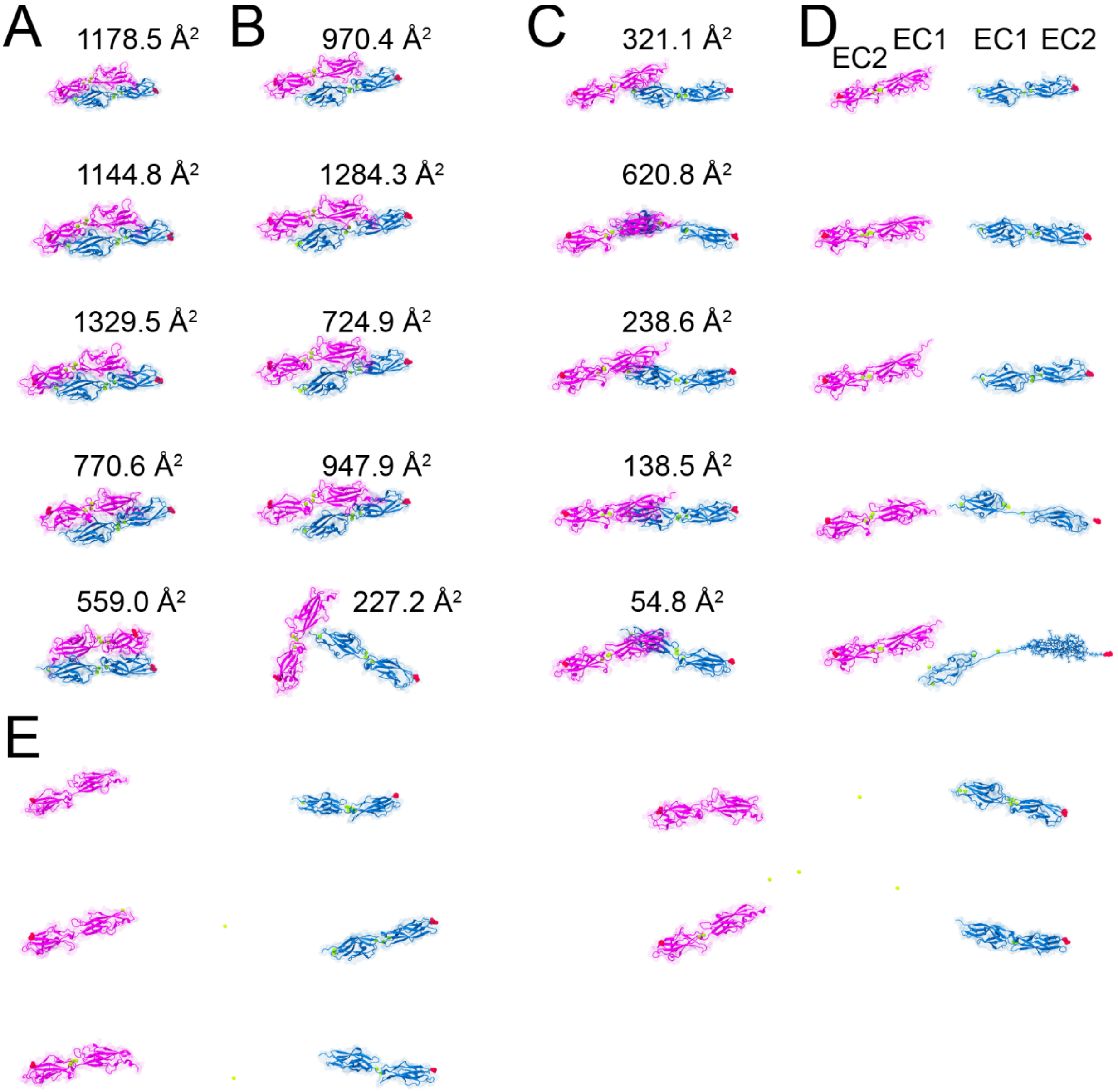

**SFigure 6: Bayesian-steered Boltz-2 predictions for the dimeric handshake formed by the CDH23 and PCDH15 tips at varying end-to-end distances.** The colvar was defined as the distance between centers of mass of the C-terminal residues of each monomer. Each prediction was generated using its own collection of 100 Feynman-Kac particles and selected by taking the resulting highest probability particle. Buried surface area (BSA) is shown in cases where it is nonzero. *A*. Unsteered Boltz-2 predictions whose colvars natively take an approximate value of 100 Å. *B-E*. Bayesian-steered Boltz-2 predictions with the colvar harmonically restrained at 120 Å, 150 Å, 250 Å, and 350 Å, respectively. Protein shown as in Figure 3D. Some panels are also shown in Figure 3.

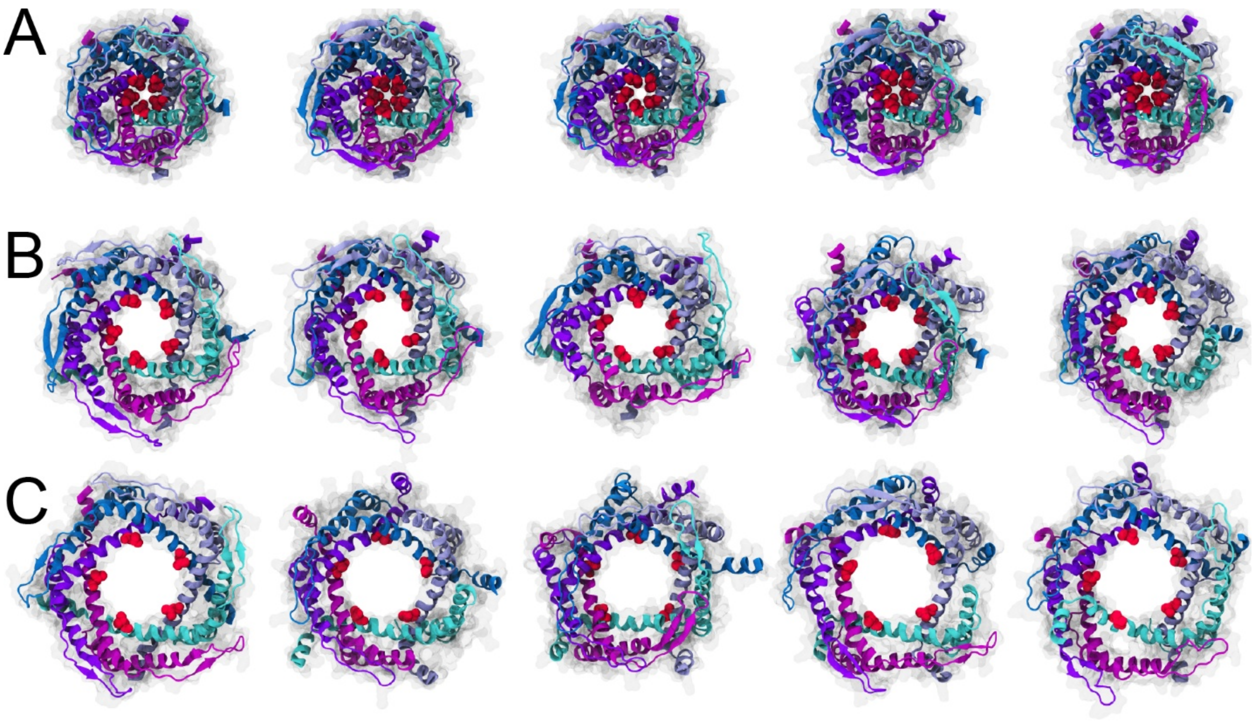

**SFigure 7: Bayesian-steered Boltz-2 predictions for MscL with top-down view of the pore at varying values of the pore colvar.** The colvar was defined as a vector whose five components were the distances of the centers of mass of each chain's valine 21 residue from the common center of mass of all five residues. Each prediction was generated using its own collection of 100 Feynman-Kac particles and selected by taking the resulting highest probability particle. *A*. Unsteered Boltz-2 predictions whose colvars natively take an approximate value of 5 Å. *B*-*C*. Bayesian-steered Boltz-2 predictions with the colvar harmonically restrained at 10 Å and 15 Å respectively. Protein shown as in Figure 4. Cytoplasmic domain is omitted for visualization purposes (see SFigure 9). Some panels are also shown in Figure 4.

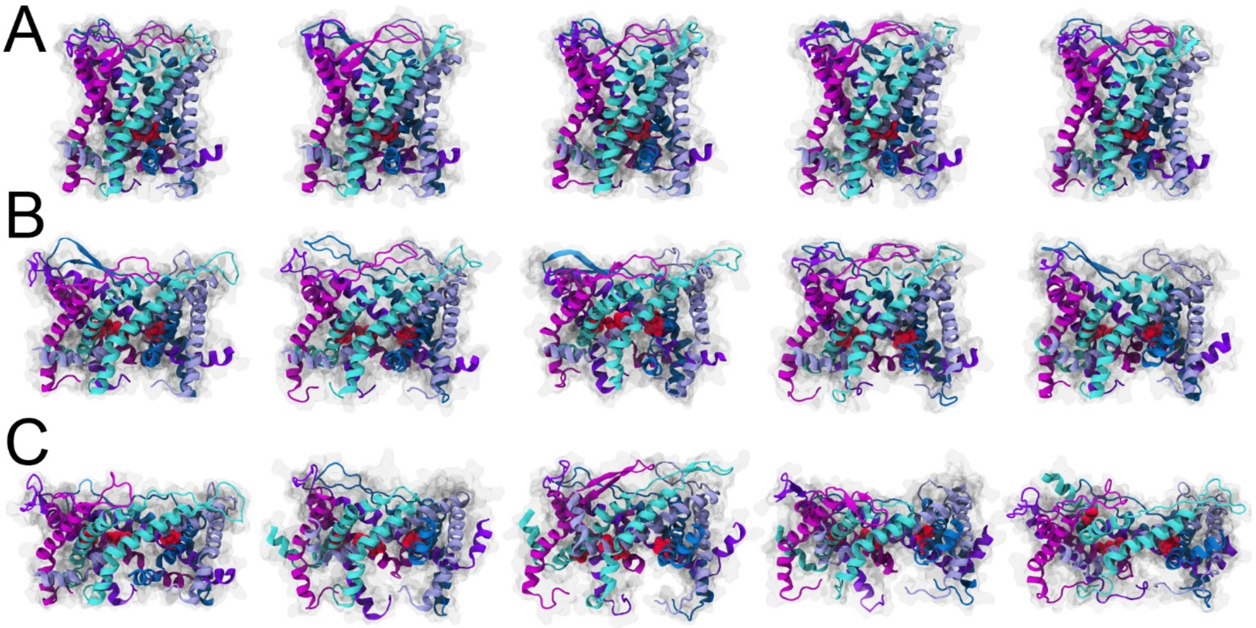

**SFigure 8: Bayesian-steered Boltz-2 Predictions for MscL with side view of the pore at varying values of the pore colvar.** The colvar was defined as a vector whose five components were the distances of the centers of mass of each chain's valine 21 residue from the common center of mass of all five residues. Each prediction was generated using its own collection of 100 Feynman-Kac particles and selected by taking the resulting highest probability particle. *A*. Unsteered Boltz-2 predictions whose colvars natively take an approximate value of 5 Å. *B-C*. Bayesian-steered Boltz-2 predictions with the colvar harmonically restrained at 10 Å and 15 Å respectively. Protein shown as in Figure 4. Cytoplasmic domain is omitted for visualization purposes (see SFigure 9). Some panels are also shown in Figure 4.

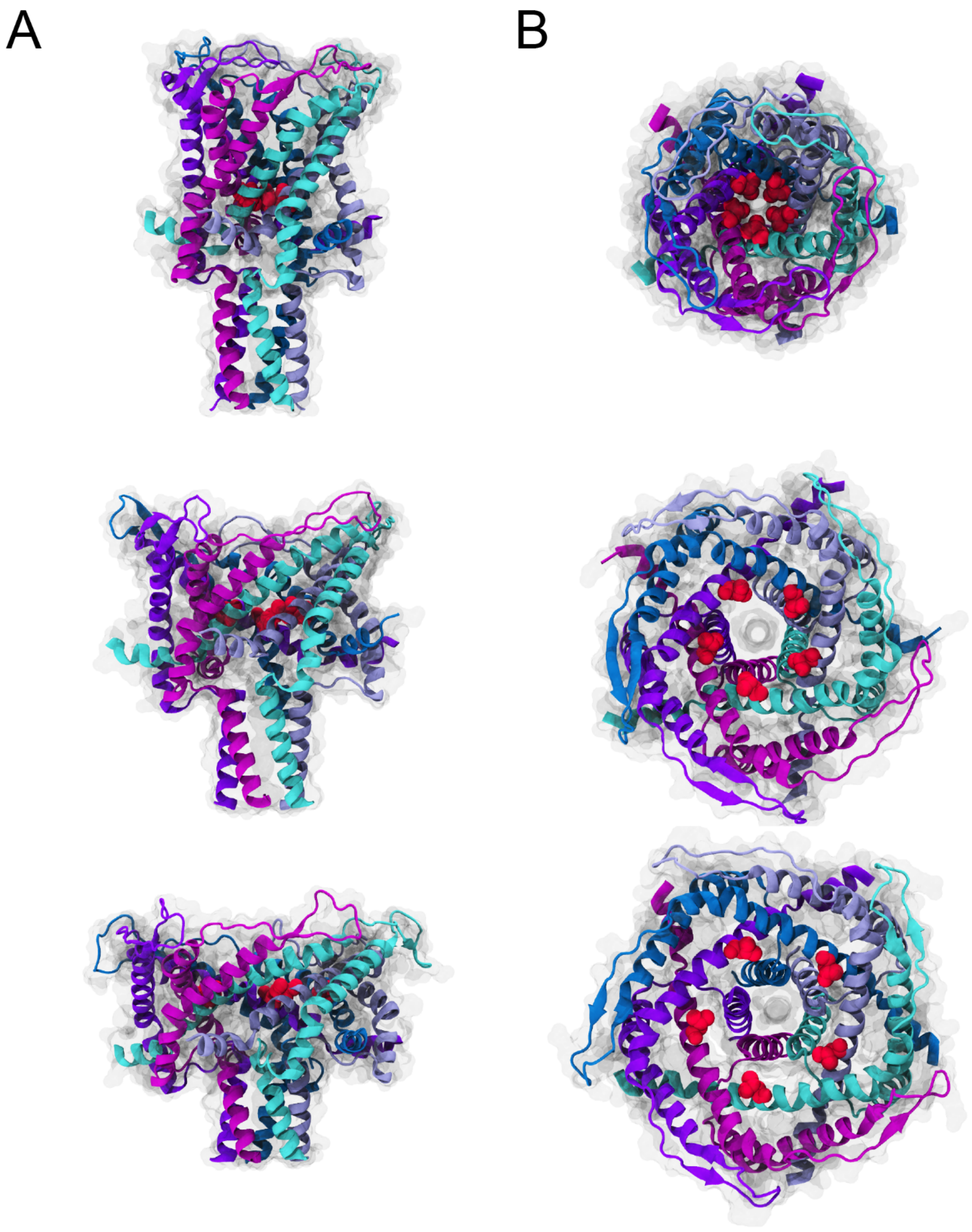

**SFigure 9: Representative Bayesian-steered Boltz-2 predictions for MscL showing its intracellular domain at varying values of the pore colvar.** The colvar was defined as a vector whose five components were the distances of the centers of mass of each chain's valine 21 residue from the common center of mass of all five residues. Each component of the colvar was unbiased and harmonically restrained at 10 Å and 15 Å. *A*. Side view. *B*. Top-down view. The pore opened in the transmembrane region while the intracellular domain stayed closed. Protein shown as in Figure 4.

*STable 1:* DNA and protein sequences used.

| |
|---|
| **DNA**: |
| GGGCGGCGACCT |
| AGGTCGCCGCCC |
| **Titin I91**: |
| SSLIEVEKPLYGVEVFVGETAHFEIELSEPDVHGQWKLKGQPLTASPDCEIIEDGKKHILILHNCQL<br>GMTGEVSFQAANAKSAANLKVKEL |
| **PCDH15 EC8-EC10:** |
| SPVFTNSTYTVLVEENLPAGTTILQIEAKDVDLGANVSYRIRSPEVKHFFALHPFTGELSLLRSLDYE<br>AFPDQEASITFLVEAFDIYGTMPPGIATVTVIVKDMNDYPPVFSKRIYKGMVAPDAVKGTPITTVYAE<br>DADPPGLPASRVRYRVDDVQFPYPASIFEVEEDSGRVITRVNLNEEPTTIFKLVVVAFDDGEPVMS<br>SSATVKILVLHPGEIPRFTQEEYRPPPVSELATKGTMVGVISAAAINQSIVYSIVSGNEEDTFGINNIT<br>GVIYVNGPLDYETRTSYVLRVQADSLEVVLANLRVPSKSNTAKVYIEIQDE |
| **PCDH15 EC9-MAD12**: |
| NDYPPVFSKRIYKGMVAPDAVKGTPITTVYAEDADPPGMPASRVRYRVDDVQFPYPASIFDVEED<br>SGRVVTRVNLNEEPTTIFKLVVVAFDDGEPVMSSSATVRILVLHPGEIPRFTQEEYRPPPVSELAAR<br>GTVVGVISAAAINQSIVYSIVAGNEEDKFGINNVTGVIYVNSPLDYETRTSYVLRVQADSLEVVLAN<br>LRVPSKSNTAKVYIEIQDENDHPPVFQKKFYIGGVSEDARMFASVLRVKATDRDTGNYSAMAYRLI<br>IPPIKEGKEGFVVETYTGLIKTAMLFHNMRRSYFKFQVIATDDYGKGLSGKADVLVSVVNQLDMQ<br>VIVSNVPPTLVEKKIEDLTEILDRYVQEQIPGAKVVVESIGARRHGDAYSLEDYSKCDLTVYAIDPQT<br>NRAIDRNELFKFLDGKLLDINKDFQPYYGEGGRILEIRTPEAVTSIKKRG |
| **CDH23 EC1-2**: |
| QVNRLPFFTNHFFDTYLLISEDTPVGSSVTQLLARDMDNDPLVFGVSGEEASRFFAVEPDTGVV<br>WLRQPLDRETKSEFTVEFSVSDHQGVITRKVNIQVGDVNDNAPTFHNQPYSVRIPENTPVGTPIFI<br>VNATDPDLGAGGSVLYSFQPPSPFFAIDSARGIVTVIQELDYEVTQAYQLTVNATDQDKTRPLSTLA<br>NLAIIITD |
| **PCDH15 EC1-2**: |
| QYDDDWQYEDCKLARGGPPATIVAIDEESRNGTILVDNMLIKGTAGGPDPTIELSLKDNVDYWVL<br>LDPVKQMLFLNSTGRVLDRDPPMNIHSIVVQVQCVNKKVGTVIYHEVRIVVRDRNDNSPTFKHE<br>SYYATVNELTPVGTTIFTGFSGDNGATDIDDGPNGQIEYVIQYNPEDPTSNDTFEIPLMLTGNVVLR<br>KRLNYEDKTRYYVIIQANDRAQNLNERRTTTTTLTVD |
| **MscL**: |
| MLKGFKEFLARGNIVDLAVAVVIGTAFTALVTKFTDSIITPLINRIGVNAQSDVGILRIGIGGGQTIDL<br>NVLLSAAINFFLIAFAVYFLVVLPYNTLRKKGEVEQPGDTQVVLLTEIRDLLAQTNGDSPGRHGGR<br>GTPSPTDGPRASTESQ |
